# Aerobic glycolysis and lactate regulate histone H3K18Lactylation occupancy to fine-tune gene expression in developing and mature retina

**DOI:** 10.1101/2025.04.17.649437

**Authors:** Mohita Gaur, Xulong Liang, Matthew J. Brooks, Ke Jiang, Anjani Kumari, Milton A. English, Paolo Cifani, Maria C. Panepinto, Jacob Nellissery, Robert N. Fariss, Laura Campello, Claire Marchal, Anand Swaroop

## Abstract

High aerobic glycolysis in retinal photoreceptors, as in cancer cells, is implicated in mitigating energy and metabolic demands. Lactate, a product of glycolysis, plays a key role in epigenetic regulation through histone lactylation in cancer. Here, we demonstrate that increased ATP production during retinal development is achieved primarily through augmented glycolysis. Histone lactylation, especially H3K18La, parallels enhanced glycolysis and lactate in developing retina and in retinal explants. Multi-omics analyses, combined with confocal imaging, reveal the localization of H3K18La near H3K27Ac in euchromatin at promoters of active retinal genes. H3K18La and gene expression are also concordant with glucose metabolism in retinal explants. Evaluation of accessible chromatin at H3K18La marked promoters uncovers an enrichment of GC-rich motifs for transcription factors of SP, KMT and KLF families, among others, indicating specificity of H3K18La-mediated gene regulation. Our results highlight glycolysis/lactate/H3K18La as a regulatory axis in fine-tuning gene expression in developing and mature retina.

## Introduction

Augmented glycolysis, even in the presence of oxygen (aerobic glycolysis), and consequently high lactate production are key hallmarks of cancer cells ^1^. The Warburg effect not only yields sufficient energy for proliferation but also impacts multiple anabolic and signaling pathways ^2^. Key glycolytic enzymes and their products, primarily lactate, participate in regulatory steps that favor cell proliferation and tumorigenesis ^3,4^. In addition to its direct linkage to mitochondrial metabolism, the lactate serves as a critical modifier of distinct cellular activities including DNA damage repair via lactylation of XRCC1 protein ^5^ and gene regulation by lactylation of histones in tumors ^6^. Lactylation of histones as well as other histone marks serve as docking sites for chromatin modifiers and transcription factors (TFs), linking the metabolic state of cells to transcriptional output ^7,8^. The studies on metabolism-epigenome relationship have largely focused on stem cells and cancer, which have unique metabolic demands for continuous proliferation and rapid biomass duplication.

As in cancer, Warburg also observed high aerobic glycolysis in the retina ^1^, a highly metabolic yet terminally differentiated non-proliferative tissue. The vertebrate retina consists of an array of diverse and highly specialized neurons that are stratified in three cellular layers for photon capture and processing of visual information ^9^. The rod and cone photoreceptors account for almost 75% of the cells in most mammalian retinas and initiate the process of vision by converting photons to electrical signals through the process of phototransduction ^10^. Neurons in the inner retina layers, including divergent subtypes of bipolar, horizontal, amacrine and ganglion cells, subsequently process the visual signals for transmission to brain via the optic nerve ^9^. The human retina includes over 105 million densely packed rods and about 6 million cones that permit vision in dim- and daylight, respectively, by closing the cGMP-gated channels in response to photon capture ^10^. Maintenance of photoreceptor physiological state and daily renewal of the membranous discs at the tip of outer segment place enormous energy and anabolic demands on the retina ^11–15^.

Glucose is the energy source in the neural retina, with photoreceptors being the primary consumer ^16^. Aerobic glycolysis in photoreceptors is implicated in maintaining high anabolic activity and survival ^17,18^. Glucose uptake in photoreceptors depends on the retinal pigment epithelium (RPE), which serves as a barrier to the choroidal blood vessels while concurrently ensuring a steady supply of glucose, oxygen and nutrients. The excess lactate produced by glycolysis in photoreceptors is transported to the RPE, serving as the substrate for oxidative phosphorylation (OXPHOS) and thereby suppressing glucose utilization by RPE ^19^. This complementary relationship between photoreceptors and RPE is suggested to create a metabolic ecosystem ^20–22^. Nonetheless, we currently have poor understanding of how the retina adjusts to changes in energy and metabolic needs during development, aging and disease conditions.

The advent of global ‘omics’ studies have begun to unravel the interdependence of epigenome modifications and metabolic pathways ^8,23^. Metabolic intermediates produced primarily from glycolysis and OXPHOS serve as key substrates for multitude of histone modifications ^24,25^; these include acetyl-CoA and acyl Co-A derivatives that modify lysine residues on histone proteins ^26,27^. In addition, epigenetic role of the lactyl-CoA, a derivative of lactate produced in glycolysis, has been identified as a modifier of lysine in histones (KLa) ^6^. In cancer cells, histone KLa acts as a key sensor of metabolic changes, translating into stable gene expression patterns to promote tumor progression^28^. The extensively studied lactylation site on histone, H3K18La, is recognized as a tissue specific active mark and exhibits similarities to histone acetylation (H3K27Ac) ^29^. Furthermore, roles of histone KLa in cell fate determination, neurogenesis, and hair cell regeneration in cochlea ^30–33^ further emphasize its significance in proliferating cells that exhibit heightened glycolytic activity. However, comprehensive functions of high lactate and lactylation in post-mitotic yet glucose-dependent retinal cells are poorly understood.

The lactate levels in the mammalian retina are 5 to 10 times higher than other organs in the body ^34^. Specifically, the retinal photoreceptors exhibit prominent expression of lactate dehydrogenase A (LDHA) which converts pyruvate to lactate ^15^. More recently, glycolysis and enhanced lactate levels have been implicated in mediating morphogenesis as well as transcriptional patterns in developing mouse eye organoids^35^. Genetic alterations that affect glycolysis can influence the structure of the photoreceptor outer segment ^17^. Additionally, metabolic underpinnings of photoreceptor dysfunction are correlated to glucose uptake and metabolism in retinal degenerative disorders ^18,36^. These studies point to broader regulatory functions of high lactate levels during retinal development and in maintaining tissue homeostasis.

In this report, we directly demonstrate augmented histone H3K18La by high lactate and glycolytic flux during development and in enhanced glucose conditions. H3K18La peaks near promoter regions partially colocalize with H3K27Ac in the euchromatin and are associated with higher expression of genes involved in retinal development and/or photoreceptor function. We also identify TF binding motifs, including those of Krüppel-like factors (KLFs) and specificity proteins (SPs) families, at accessible H3K18La peaks. Our results implicate aerobic glycolysis and lactate levels, and consequently augmented histone H3 lactylation at K18 residue, as key mediators of transcriptional program in the developing and mature retina, and especially in the photoreceptors.

## Results

### Enhanced glycolytic flux contributes to lactate accumulation in developing retina

We used Seahorse XFe24 Analyzer to measure glycolysis (extracellular acidification rate or GlycoECAR) and mitochondrial respiration (oxygen consumption rate or MitoOCR) of developing and mature mouse retina (**Figure 1A-C**). We noted that GlycoECAR continues to increase after postnatal day (P)6 (**Figure 1B**), whereas MitoOCR is only marginally higher in mature (P28) retina (**Figure 1C**). Total ATP production, contributed from both glycolysis and mitochondrial respiration, is significantly enhanced as development proceeds (**Figure 1D**). ATP production contributed by OXPHOS exhibits little change from P2 to P28. Until P10, mitochondrial respiration appears to serve as the primary source of ATP, with glycolysis being a minor contributor. However, glycolysis flux continues to increase from P6 onward and becomes the major energy source after P14, as retinal neurons terminally differentiate, form synapses and become functionally mature around P14 (**Figure 1D, E**). During retina development, quantifications using bioluminescent Lactate-Glo™ assay demonstrate a steady increase in lactate levels (**Figure 1F**). Significantly higher lactate levels were detected in retinas at late developmental stages (P14 and P28) when compared to early developmental stages (P2 and P6) and correlate with enhanced glycolysis (**see Figure 1B**). We also detect a significant increase in lactate dehydrogenase activity from P6 to P10 and P28 (**Figure 1G**). Overall, comprehensive GlycoECAR and mitoOCR measurements indicate augmented aerobic glycolysis in developing retina to meet higher energy and metabolic demands (**Figure 1H**).

**Figure 1:**
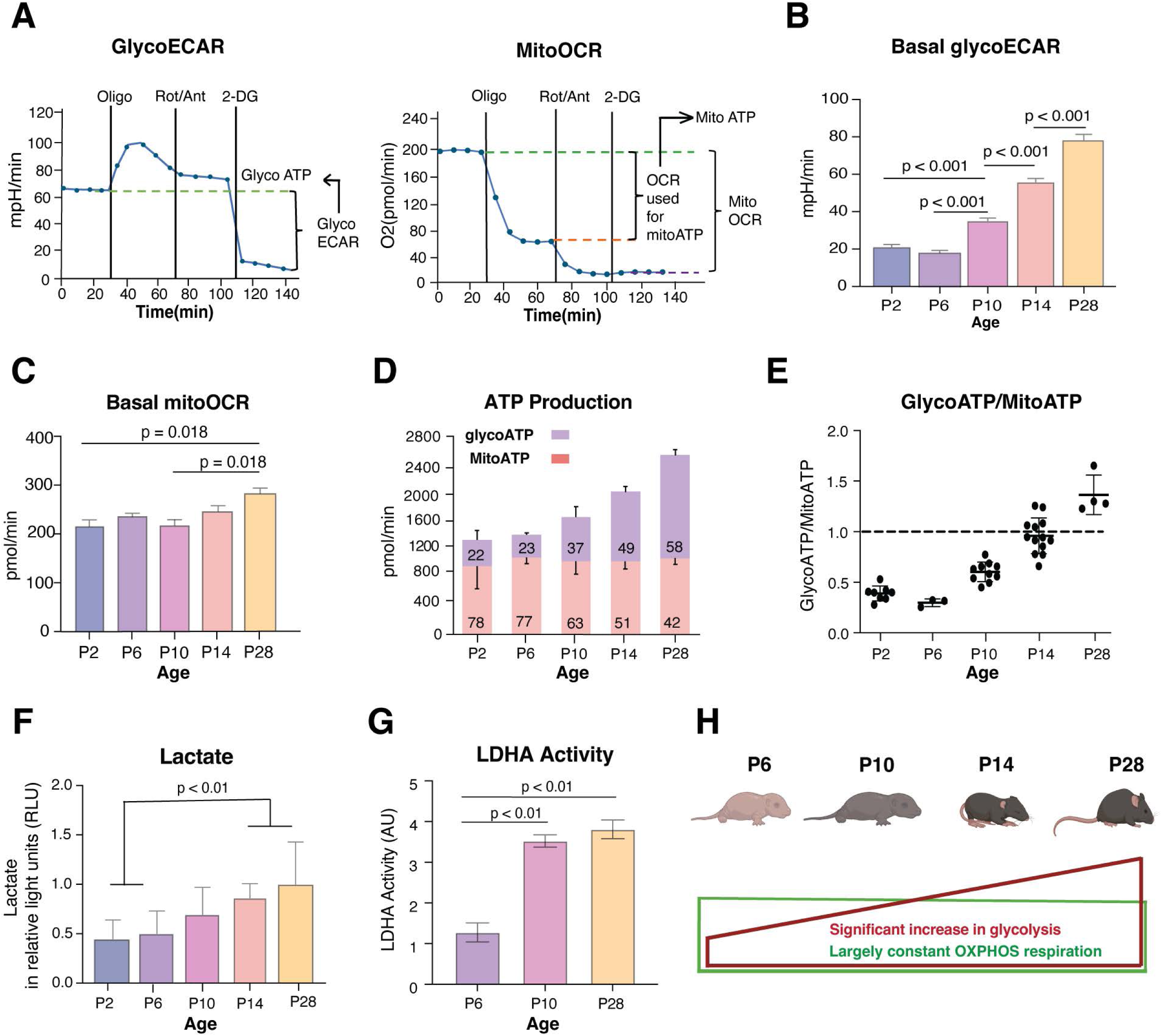
Enhanced contribution of aerobic glycolysis and lactate to ATP generation during retinal development. A) Graphical representation of glycolytic flux profile (left) and mitochondrial respiration profile (right) using Seahorse XFe24 Analyzer. The left graph represents extracellular acidification rate (ECAR), which is largely proportion to glycolytic activity of cells and the right graph represents oxygen consumption rate (OCR), which is an indicator for mitochondrial respiration. The X-axis shows duration of the assay, and the timing of sequential inhibitor additions are indicated by vertical lines. The initial baseline (0 to 36 min) represents normal glycolysis level and respiration level before any metabolic interventions. With Oligomycin (an inhibitor of mitochondrial complex V), oxygen consumption drops drastically whereas glycolytic flux displays a sharp increase followed by a slight decline. The addition of Rotenone/Antimycin A (inhibitors to complex I and III) further shuts down mitochondrial respiration but with minimal influence on glycolysis flux. The final addition of the glycolytic inhibitor 2-Deoxy-D-glucose (2-DG) completely abrogates glycolysis. B) and C) Basal GlycoECAR and basal mitoOCR were quantified from Seahorse assays using retinal punches isolated from mice at different developmental age groups (n = 13– 35 retinal punches per age group from 4 mice). Data are presented to compare the metabolic activity between age groups, with glycoECAR reflecting glycolytic activity and mitoOCR reflecting mitochondrial respiration. D) ATP production rates calculated from collated MitoOCR and GlycoECAR data from B and C. The number on the bar indicates percentage. E) Ratio of glycolysis derived ATP to mitochondrial derived ATP was calculated from data collated for D. F) Lactate concentration was quantified for different age groups in retina tissue samples using Lactate-Glo Luciferase Assay, n= 4 except for P28, n = 3. Data plotted as mean luminescence values (relative light units – RLU) from each replicate per age group. G) Lactate Dehydrogenase (LDH) Assay Kit (ab102526, Abcam) was used to measure LDH activity in the mouse retina from different age groups (P6, P10, P28; n=4 per age group). H) Pictorial representation of ATP production during retinal development, showing enhanced glycolysis derived ATP production (represented as red rectangle), however proportional percentage contribution from mitochondrial derived ATP remains largely constant with age (represented as green rectangle). All data in the figures are presented as mean ± SEM. Statistical significance is indicated by p-values displayed above the bars. One-way ANOVA with Tukey’s post hoc comparison was used to determine the significance.

### H3K18La in the retina correlates with enhanced glycolysis

We hypothesized that lactate levels contribute to chromatin state and gene expression via histone lactylation (**Figure 2A**). To test this hypothesis, we first performed high-resolution mass spectrometry analysis of histones from the P28 mouse retina. KLa is abundantly present across histone proteins (**Figure 2B, Figure S1A, Table S1**). Immunoblot analysis revealed a significant increase in H3K18La levels during retina development (from P2 to P28), but no significant change was evident in Pan-lactyl and H4K12La signals (**Figure 2C**).

**Figure 2:**
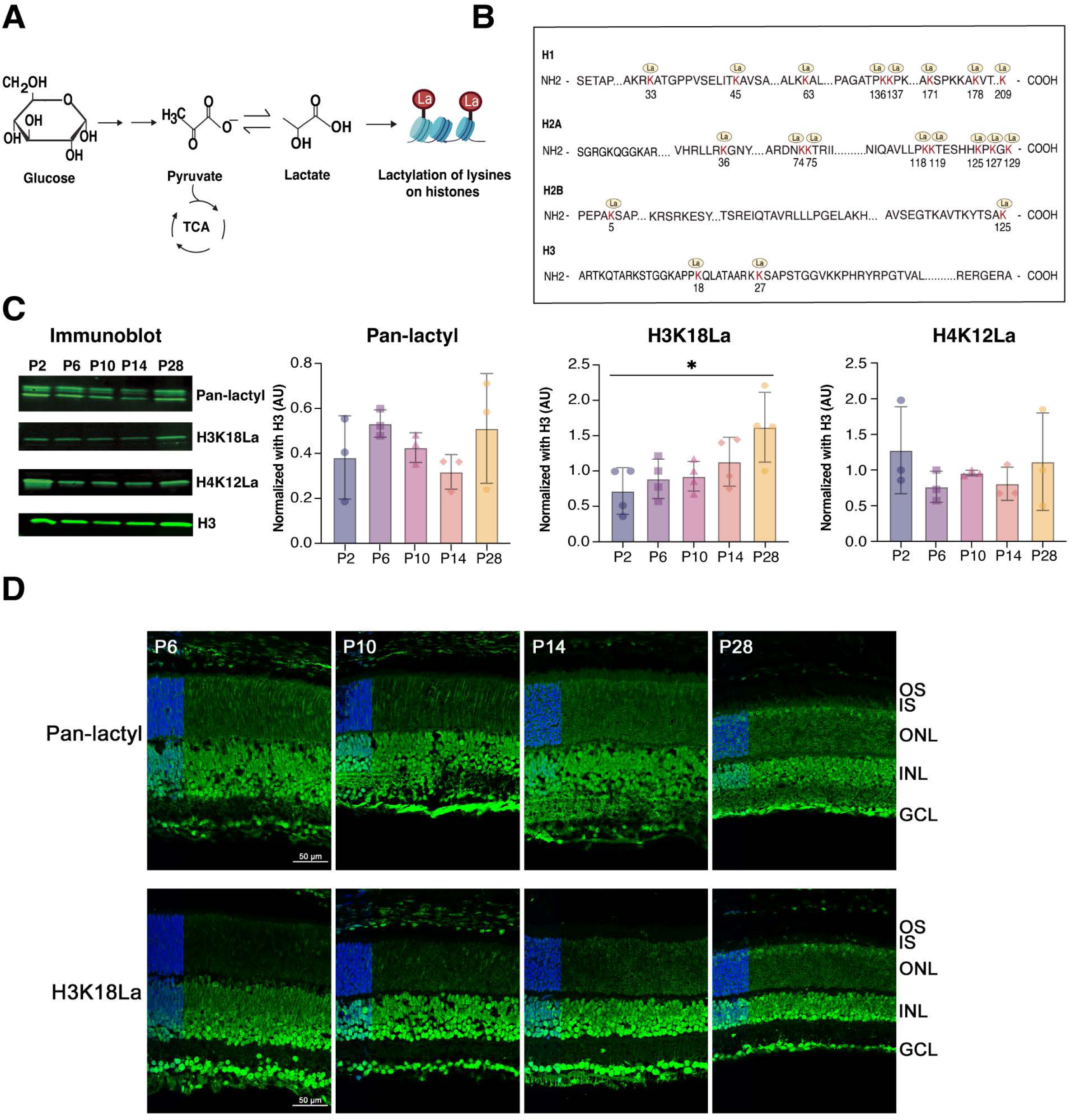
Histone lactylation in the retina. A) Graphical illustration of non-metabolic role of high lactate production in retinal photoreceptors cells because of Warburg’s effect. The lactate produced can act as chemical tag “La” for histone which further modulates gene transcription in retina. B) Illustration of histone lysine (K) lactylation (La) sites identified by mass spectrometry in derivatized tryptic-digested peptides from adult (P28) mouse retina. The identified lactylation sites on lysine residues are highlighted in red and marked by yellow circle icons with “La” written inside. See also Figure S1A for annotated HCD Spectrum and Table S1 for a complete list of identified lactylation sites, represented as carboxyethyl in the modification list. C) Immunoblot-based quantification of histone KLa in purified histone samples at different ages of mice for Pan-lactyl (n = 3), H3K18La (n=4) and H4K12La (n=3) with the experiment repeated twice. ImageStudio software was used for quantification. Statistical significance was analyzed using one-way ANOVA with post hoc Tukey’s test. Immunoblots for Pan-lactyl, H3K18La, and H4K12La show molecular weights of 15 kDa. All data are presented as mean ± SEM. *, p<0.05. D) Immunohistochemical analysis of Pan-KLa or H3K18La in retina tissue samples from different age groups of mice. Inset represents a zoomed-out view with DAPI staining in the ONL on the left. Abbreviations: OS, outer segment; IS, inner segment; ONL, outer nuclear layer; INL, inner nuclear layer; GCL, ganglion cell layer. Scale bar = 50 µm.

We then performed immunohistochemistry (IHC) of mouse retina sections using anti-H3K18La and Pan-lactyl antibodies. During early postnatal development, prominent H3K18La and Pan-lactyl immunostaining is observed in the nuclei of retinal cells as well as in axonal and dendritic processes (**Figure 2D**). However, the fluorescence appears somewhat diffused in early stages of photoreceptor differentiation and gradually acquires a more defined pattern that is consistent with unique euchromatin architecture in mature photoreceptors (**Figure 2D**). H3K18La antibodies exhibit stronger signal intensity at P28, consistent with immunoblotting. Notably, H3K18La levels closely correlate with glycolysis in developing and mature retinal photoreceptors, whereas Pan-lactyl immunostaining is broadly enhanced in the nuclei of most retinal cell types, indicating widespread lactylation (**Figure 2D**).

### Regulation of histone lactylation by glucose metabolism

To directly examine the relationship of histone lactylation with glycolysis, we performed a glucose modulation experiment in explant cultures of P28 mouse retinas. We exposed retinal explants to two different conditions: elevated glucose levels (5 mM to 25 mM) and addition of 10-20 mM 2-deoxy-D-glucose (2-DG), a non-metabolizable glucose analog that inhibits glycolysis. We then measured the lactate secreted in the media, together with Pan-lactyl, H3K18La and H3K27Ac changes in the retinal tissue (**Figure 3A**). Both lactate production and histone lactylation levels (both H3K18La and Pan-lactyl; n=3, p<0.05) are induced by glucose in a dose-dependent manner (**Figure 3B, C**); however, incubation with 2-DG decreases both lactate and histone KLa levels significantly (Pan-lactyl p<0.01; H3K18La p<0.05, n=3) (**Figure 3D, E**). Curiously, high glucose conditions don’t significantly alter H3K27Ac levels in retinal explants (**Figure 3C**); yet the treatment of explants with 2-DG, which competes with glucose for binding, significantly reduces the H3K27Ac levels (p<0.05) (**Figure 3E**). Thus, lactate levels and consequently histone lactylation are dependent on glycolytic flux in retinal explants.

**Figure 3:**
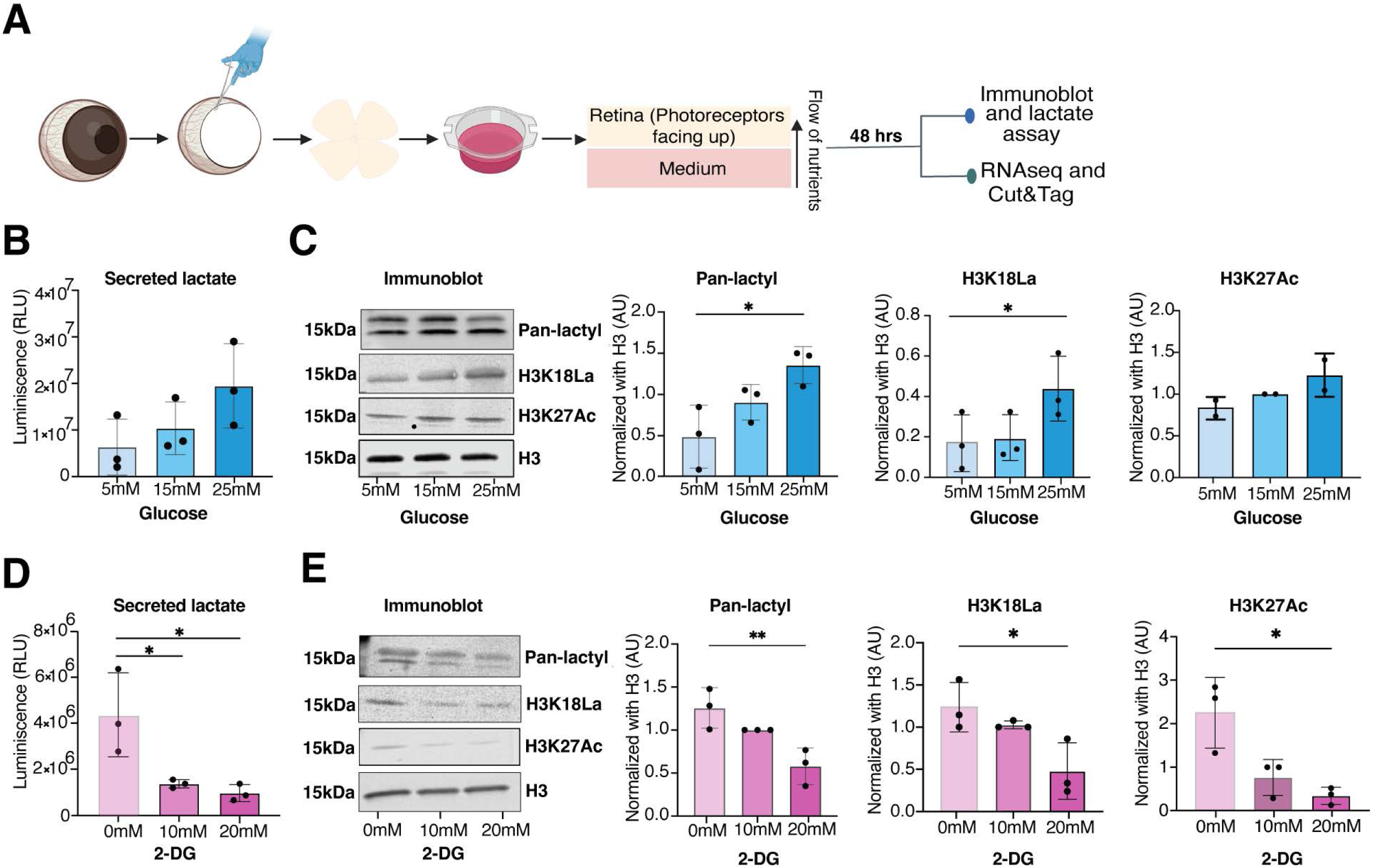
Glycolysis regulates histone lactylation in the retina. (A) Representation of the retinal explant workflow. Key steps in the experiment are shown. The isolated retina was cultured in explant media for 48 hours before proceeding for subsequent experiments as outlined in the figure. (B, D) Extracellular lactate levels using bioluminescence assay and (C, E) histone lactylation (KLa) levels using immunoblot were measured in retinal explants cultured for 48 hours in different concentrations of glucose (5 mM, 15 mM, and 25 mM) or 2-Deoxy-D-glucose (2-DG) (10 mM and 20 mM) with 5 mM glucose as a baseline control in 2-DG experiments. Extracellular lactate, the amount of lactate secreted into the culture media was quantified, and the amount of lactate was correlated with luminescence levels, expressed in relative light units (RLU) on Y axis as shown. For (C, E) immunoblots, histones were extracted from retinal tissues cultured under the same conditions for 48 hours. Pan anti-KLa, H3K18La, and H3K27Ac antibodies detected protein bands at approximately 15 kDa. Data are presented as mean ± SEM (n = 3 biological replicates, *, p<0.05; **, p<0.01). Statistical significance was determined using one-way ANOVA followed by Sidak’s multiple comparisons test.

### Chromatin context-dependent gene regulation by H3K18La

We then investigated genome-wide occupancy of H3K18La, using CUT&Tag assay, during postnatal mouse retinal development (P4, P10 and P28). Consensus peaks were determined by using q-value threshold of 1×10^-6^ and selecting those observed in at least two (out of 3) replicates. Consensus H3K18La peak numbers increase from P4 (51,085 peaks) to P28 (64,973 peaks) but show a decrease at P10 (30,777 peaks) (**Figure 4A**). Interestingly, the number of promoters with a H3K18La peak exhibit a dramatic increase in the adult retina (P28) with higher distribution proximal to TSS, even though a majority of H3K18La peaks are detected in gene bodies (**Figure 4B,C**). Peaks being called at P4 appear to be widely distributed and lower in signal intensity prior to coalescing to fewer, but more robust peaks at P10, and before finally establishing the highest peak signals at P28 (**Figure S1B**). Select examples of genes involved in lactate production (*Aldoa* and *Ldha*) and phototransduction (rhodopsin, *Rho*) demonstrate the dynamics of H3K18La binding during development (**Figure 4D**). Gene ontology (GO) enrichment analysis of genes associated with H3K18La peaks in promoters, enhancers or gene bodies shows differential enrichment of cellular pathways (**Figure S1C, Table S2-S4**). H3K18La peaks are enriched in promoters of genes involved in canonical Wnt signaling, synaptic membrane adhesion and circadian regulation, whereas those in the enhancer regions exhibit dominance of retinal functions including eye development, visual perception and axon guidance. The latter observation is consistent with a higher correlation of enhancer elements with cell-type-specific genes compared to the promoters ^37^. The H3K18La peaks in gene bodies are enriched for ERC1/2 cascade, synapse assembly, Wnt signaling and photoreceptor cell maintenance.

**Figure 4:**
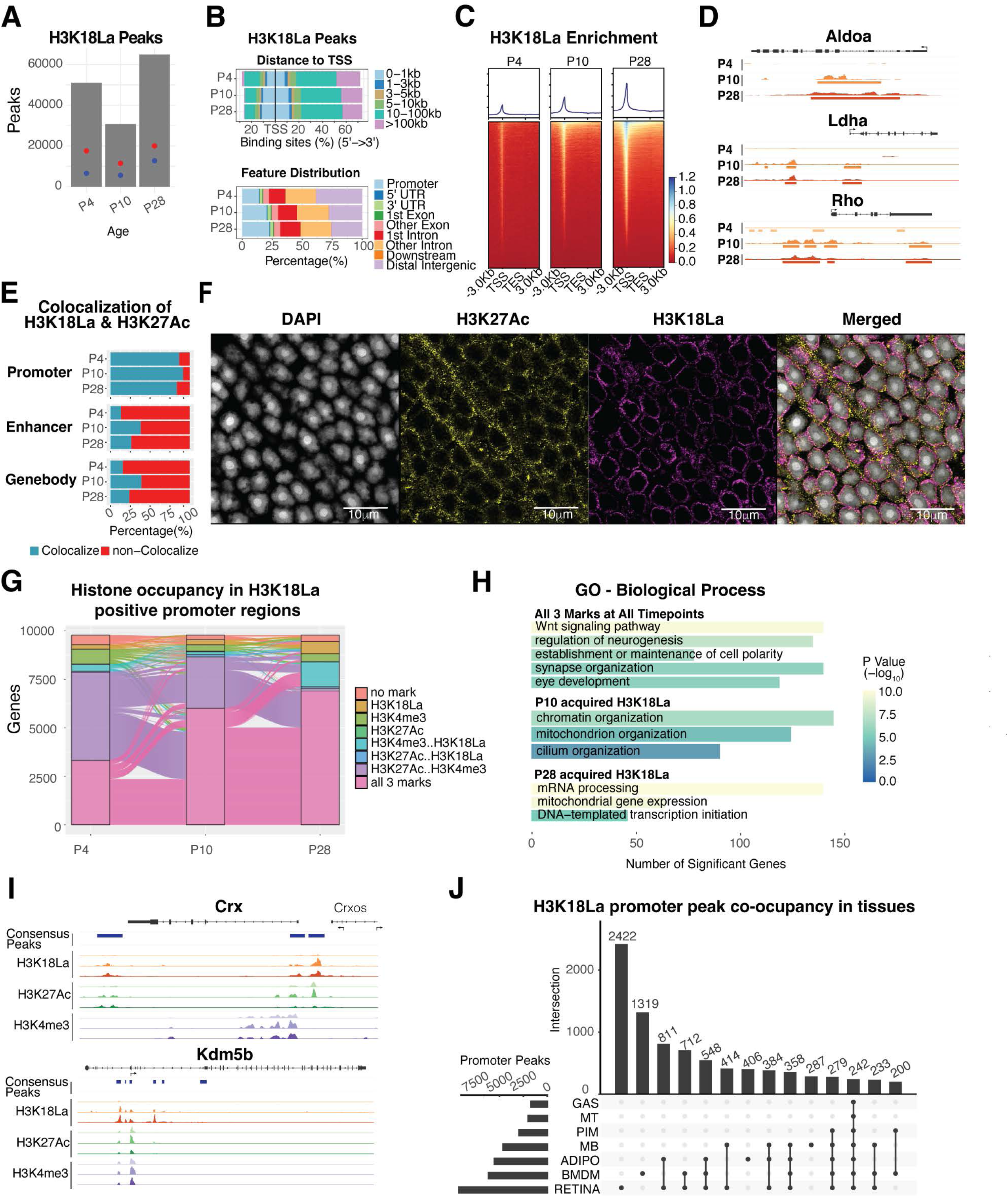
Genome-wide profiling of H3K18La. (A) The number of consensus peaks passing a 1×10^-6^ FDR for each replicate per post-natal timepoint. The red dot indicates the number of genes containing a peak (17,647, 11,511, and 20,087 respectively), whereas the blue dot represents the number of genes containing a peak in the proximal promoter (6,646, 5,627, and 12,822 respectively). (B) Upper panel: Percentage of H3K18La bound consensus peak loci distance relative to TSS. Lower panel: Percentage of H3K18La bound consensus peak loci to gene annotation feature. (C) Heatmap of H3K18La-bound peak signal enrichment for all genes and their flanking 3kb region. (D) Genomic histogram traces of H3K18La at each timepoint for genes involved in glycolysis (*Aldoa* and *Ldha*) and the phototransduction cascade in rod photoreceptors (*Rho*). The histogram traces are group scaled for each gene. The bars under each histogram represent consensus peaks. (E) Colocalization of H3K18La peaks with H3K27Ac ^38^ bound regions. (F) Confocal immunofluorescent images showing colocalization of H3K27Ac and H3K18La in nuclear periphery of photoceptor cells. (G) Dynamic active histone mark (H3K4me3 and H3K27Ac ^38^) co-occupancy with H3K18La in the proximal promoter of genes containing H3K18La during development. (H) Selection of GO Biological Process gene sets enriched for genes from panel (G) containing all three histone marks at all timepoints or genes that acquired H3K18La loci at P10 or P28. (I) Genomic histogram traces of histone marks during development for representative selected genes found in panel H. H3K18La marks at P4, P10, and P28 (Red), H3K27Ac ^38^ at P3, P10, P21 (Green), and H3K4me3 at P3, P10, and P21 (Purple). (J) The UpSet plot illustrates the differential H3K18La co-occupancy at promoter regions of protein coding genes between retina and other metabolically distinct tissues^29^. See also Figure S2 Abbreviations: TSS, transcriptional start site; TES, transcriptional end site; GO, gene ontology; GAS, gastrocnemius; MT, post-mitotic end-state myotubes; PIM, post-ischemia macrophages; MB, myoblasts; ADIPO, adipose tissues; BMDM, bone marrow-derived macrophages.

We further evaluated the chromatin annotation of the peaks using a ChromHMM model derived from previously published retina data ^38^ (**Figure S2A**). H3K18La marks show an enrichment for active promoter annotations with strong TSS-proximal enhancer annotations becoming more prominent at P10 and P28. H3K18La peaks at enhancers and gene bodies show similar patterns of chromatin annotations, including enhancer and transcription permissive chromatin.

To gain further insights, we performed colocalization of H3K18La peaks with published retina ATAC-seq, H3K27Ac ^38^ and CUT&RUN genome occupancy data for the key photoreceptor transcription factor NRL ^39^ that is essential for rod differentiation^40^. H3K18La peaks exhibit high co-localization percentage with H3K27Ac and accessible chromatin at promoters but much lower at gene bodies or enhancers (**Figure 4E**, **Figure S2B)**. Confocal microscopy using specific antibodies validates partial colocalization of H3K27Ac and H3K18La signals in the euchromatin region at the periphery of photoreceptor nuclei (**Figure 4F**). Little or no colocalization is detected between H3K18La and H3K27me3 or NRL immunolabeling (**Figure S2C**).

Inspection of the promoters of protein coding genes containing H3K18La marks during any point in development (**Figure 4G**) reveals co-occurrence of H3K18La with previously reported active histone marks, H3K27Ac and/or H3K4me3 ^38^. Notably, co-occurrence of these marks at promoters is only 38% at P4 (early development) but increases to 84.4% in P28 (mature) retina. Predictably, the corresponding genes having all three active promoter marks reflect an enrichment of biological pathways relevant to retinal development (**Figure 4H,I, Figure S2D, Table S5**); such as, Wnt signaling, neurogenesis and eye development (*Crx* and *Gsk3b*). Acquired H3K18La peaks at P10 enrich for pathways involved in chromatin (*Kdm5b*), mitochondria and cilium organization (*Bbs4*) and at P28 transcription initiation (*Tbp*), mRNA processing and mitochondrial gene expression (*Taco1*).

We then evaluated the co-occurrence of H3K18La binding in retina with other tissues of differing metabolic activity ^29^. After processing the data in an identical manner, we tallied and compared the presence of H3K18La peaks in promoters of protein coding genes **(Figure 4J, Figure S2E)**. Notably, approximately one third of peaks are specific to the retina and indicate significant enrichment of retina-specific genes. The top enriched pathways are sensory perception and modulation of chemical synaptic transmission **(Table S6)**.

### H3K18La couples the metabolic state with gene regulation in developing retina

To understand the relationship of H3K18La at promoter regions with gene expression, we quantified 108,253 merged consensus peaks for the three ages (P4, P10 and P28) and visualized the data by Principal Component Analysis (PCA) (**Figure 5A, upper panel**) together with the published retinal gene expression data ^41^ (**Figure 5A, lower panel**). PC1 demonstrates the largest variance in the data, which can be attributed to the age. At P4 and P28, the replicates are well clustered revealing biological differences between the two age groups. In contrast, the P10 data show variability among replicates, likely reflecting the dynamic acquisition of H3K18La during an active differentiation state of morphogenesis and synapse formation. We then performed differential binding (DB) analysis of the merged and quantified consensus peaks; of these, 34,794 show significant DB (FDR < 0.01) during development **(Table S7)**. Furthermore, only 126 of the 5629 DB peaks in promoters of 5170 protein-coding genes (**Figure 5B)** exhibit a decrease in binding from P4 to P28. Pearson correlation analysis of 5629 significant DB peaks in promoters of protein-coding genes uncovers 1734 genes that demonstrate a positive correlation of > 0.8 between H3K18La DB peak quantitation and RNAseq data from developing mouse retina (**Figure 5C, left panel**), and 1253 genes revealing a negative correlation < −0.8 (**Figure 5C, right panel, Figure S2F**). Enrichment analysis of the positively correlated genes broadly reveals the pathways involved in phototransduction, synapse, and glycolysis (**Figure S2F, Table S8**). Thus, H3K18La at promoters seem to have distinct roles depending upon the chromatin context.

**Figure 5:**
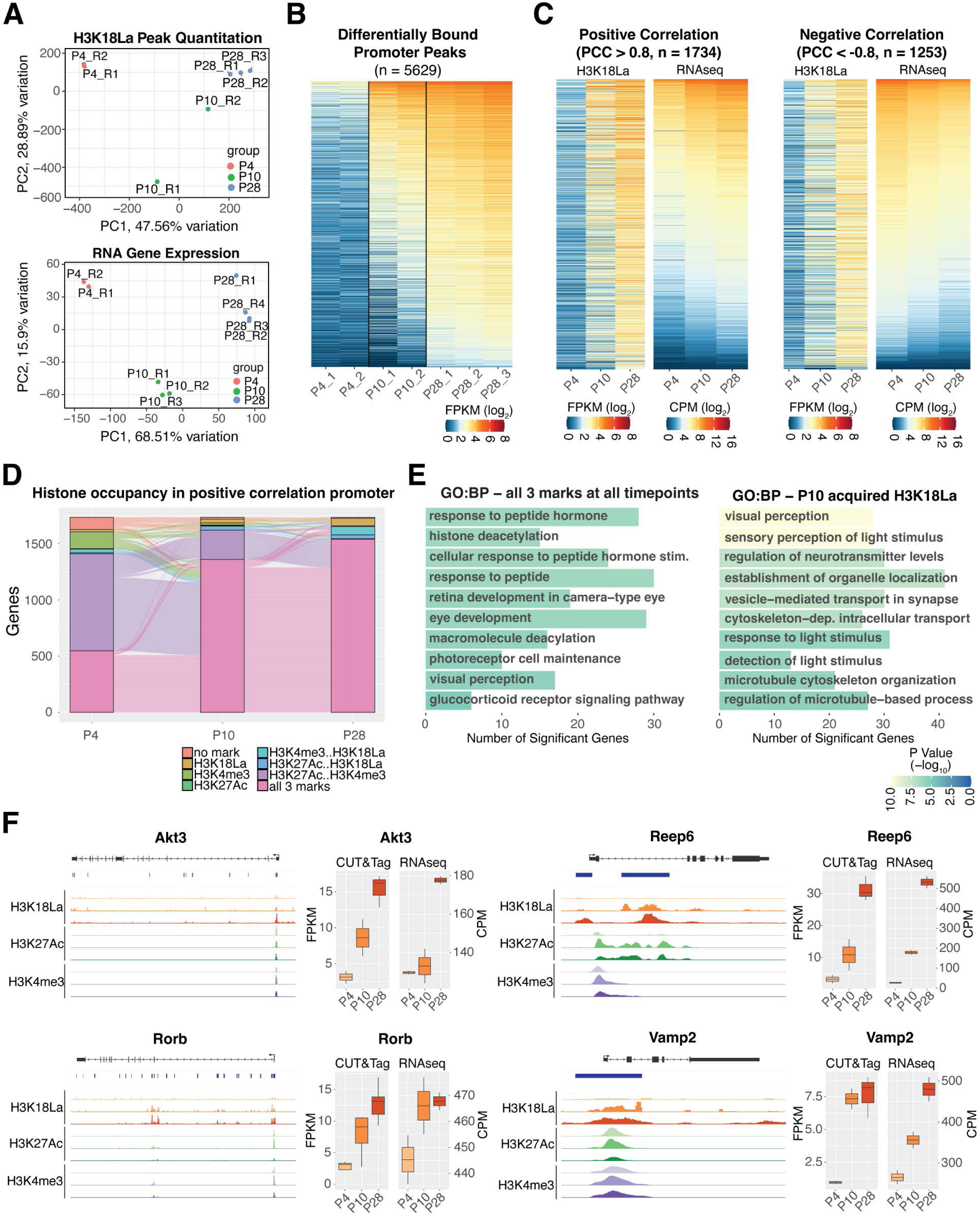
Differential binding of H3K18La over development. (A) Upper panel: Principal component analysis of quantitative peak binding for all H3K18La samples. Lower panel: Principal component analysis of RNA expression ^41^ for the same genes having H3K18La peaks. (B) Heatmap of H3K18La quantitative binding for differentially bound peaks in the promoter region of protein coding genes. (C) Left panel: Positive correlation of quantitative peaks of H3K18La found in panel (B) with retina RNA-seq expression ^41^. Right panel: Negative correlation of quantitative peaks of H3K18La found in panel (B) with retina RNA-seq expression ^41^. (D) Dynamic active histone mark (H3K4me3 and H3K27Ac ^38^) co-occupancy with H3K18La in the positive correlation set of genes from panel (C). (E) Selection of GO Biological Process gene sets enriched for genes from panel (D) containing all three histone marks at all timepoints or genes that acquired H3K18La loci at P10. (F) Left: Genomic histogram traces of histone marks during development for representative selected genes found in panel (E). H3K18La marks at P4, P10, and P28 (Red), H3K27Ac at P3, P10, P21 (Green), and H3K4me3 at P3, P10, and P21 (Purple). Right: Boxplot of quantitative binding and RNA-seq expression and CUT&Tag analysis. Abbreviations: PCA, principal component analysis; PC1, principal component 1; PC2, principal component 2; Pearson correlation coefficient (PCC); FPKM, fragments per kilobase per million reads; CPM; counts per million; GO, gene ontology; BP, biological process

We then focused on DB promoters from 1734 positively correlated genes to explore H3K18La in a broader chromatin context. Most of these promoters (57.5%) possess active histone marks, H3K27Ac or H3K4me3, at P4 prior to acquiring H3K18La and exhibit an increase in gene expression during development (**Figure 5D**). At P4, only 34.6% of positively correlated promoters have all three modifications (H3K27Ac, H3K4me3, H3K18La), yet 83.9% carry all three marks by P28 (**Figure 5D**). These genes are enriched for response to peptide hormone, histone deacetylation, and eye development (**Figure 5E, left panel, Table S9**) and include *Akt3* and *Rorb*, which contain all three histone marks from P4 onwards. *Akt3* (**Figure F, top left panel**) is important for regulating the canonical insulin/ATK/mTOR pathway and reprogramming photoreceptor metabolism and survival ^42^. *Rorb* (**Figure F, bottom left panel**) is a nuclear receptor involved in rod photoreceptor development ^43^. A robust H3K18La mark is observed at P10 in genes associated with photoreceptor maturation including visual perception and synapse processes (**Figure 5E, right panel**). As an example, *Reep6* (**Figure F, top right panel**) plays a critical role in trafficking guanylate cyclases in rod photoreceptors, and its loss results in retinopathies ^44,45^. Neuronal synaptobrevin (*Vamp2*) (**Figure 5F, bottom right panel**) facilitates fusion of synaptic vesicles ^46^. Thus, H3K18La seems to be associated with establishment of specialized neuronal functions by activating transcription from promoters of genes with a permissive chromatin context. The presence of H3K18La at P10 also indicates enhanced glycolytic activity and attainment of glucose-dependent processes necessary for effective visual function.

### Lactate promotes enrichment of H3K18La

To test whether lactate can directly influence gene expression as reported in retinal organoids ^35^, we performed CUT&Tag analyses for H3K18La and H3K27Ac together with RNA-seq in retinal explants under 5 mM and 25 mM glucose conditions (and consequently resulting in distinct lactate levels; see Fig. 3). As predicted, we observe a higher number of H3K18La peaks and peaks within promoters at 25 mM compared to 5 mM (**Figure 6A**), but with relatively little change in the total number of marks (i.e., 27,663 in 5 mM compared to 27,183 in 25 mM). For H3K27Ac, the total number of peaks (24,921 and 17,273 in 5 mM and 25 mM, respectively) and the peaks detected within genes and their promoters (12,918 and 11,369 in 5 mM and 25 mM, respectively) decrease under 25 mM glucose conditions. Furthermore, the number of peaks and their location near transcription start site (TSS) increase with 25 mM glucose condition for both histone marks (**Figure 6B, C**). This paradox for H3K27Ac of having peaks in fewer unique promoters but more peaks in promoters in general, indicates an increased number of peaks per promoter with higher glucose concentration. Visual inspection of the H3K18La and H3K27Ac peaks and their read pileups (data not shown) indicates an overall increase in signal intensity, presence of additional peaks, and merging of smaller peaks into larger robust ones with glucose supplementation (i.e., at 25 mM). PCA of the quantified peaks under 5 and 25 mM glucose conditions reveals a larger separation of peaks for H3K18La compared to what is seen for H3K27Ac (**Figure 6D**). Promoters of protein coding genes gaining H3K18La during development gain both H3K18La and H3K27Ac marks under high glucose **(Fig 6E)**.

**Figure 6:**
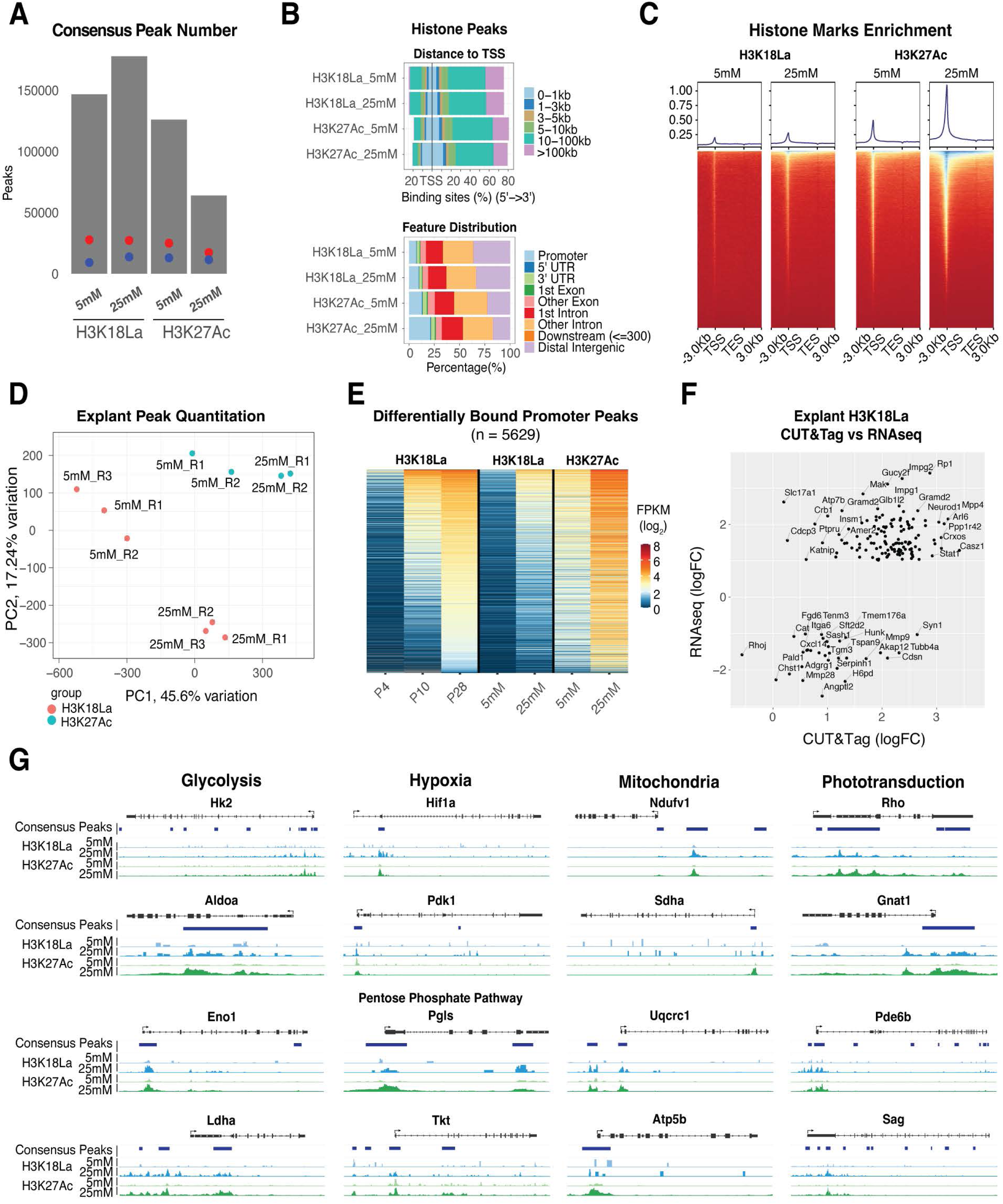
Genome-wide profiling of H3K18La in retinal explant cultures with different glucose concentration. (A) The number of consensus peaks passing a 1×10^-6^ FDR for each replicate per explant glucose concentration and histone mark. The red dot indicates the number of genes containing a peak, whereas the blue dot represents the number of genes containing a peak in the proximal promoter. (B) Upper panel: Percentage of H3K18La and H3K27Ac bound consensus peak loci distance relative to TSS. Lower panel: Percentage of H3K18La bound consensus peak loci to gene annotation feature. (C) Heatmap of H3K18La-bound and H3K27Ac-bound peak signal enrichment relative to distance from TSS/TES for all genes and their flanking 3kb region. (D) Principal component analysis of quantitative peak binding for all H3K18La and H3K27Ac samples. (E) Heatmap of peaks found from time course H3K18La differential binding analysis (Figure 5B) plotted with peak quantitation from explant CUT&Tag analysis of H3K18La and H3K27Ac. (F) Comparison of significantly, differentially expressed genes from RNA-seq explant analysis (25 mM versus 5 mM) with explant CUT&Tag differential binding results. (G) Genomic histogram traces of explant histone marks for representative genes of several affected pathways observed. The histogram traces are group normalized for each gene. The bars under each gene track represent consensus peaks. Abbreviations: TSS, transcription start site; TES, transcription end site; PCA, principal component analysis; PC1, principal component 1; PC2, principal component 2; CPM, counts per million; DB, differential bound; FPKM, fragments per kilobase per million reads; logFC, log fold-change.

RNA-seq of explant retinas were then compared with corresponding changes in H3K18La peaks. A majority of the significantly differentially expressed (FC > 2, FDR < 1%) genes show an increase in expression with higher glucose **(Fig 6F, Table S10)**; these include genes involved in retina development (*Neurod1*, *Casz1*, and *Crxos*) or those associated with retinal disease (*Impg1/2*, *Gucy2f*, *Mpp4*, *Arl6*, and *Rp1*). Additionally, genes involved in metabolic pathways and rod phototransduction demonstrate an increase in H3K18La and H3K27Ac marks with high glucose **(Fig 6G)**. Taken together, we conclude a linkage between dynamic changes in H3K18La and gene expression.

### Accessible chromatin regions with H3K18La are enriched for GC rich motifs

To further explore the mechanism by which H3K18La impacts gene expression, we evaluated accessible chromatin regions containing H3K18La for TF binding motifs (**Figure S3A**). We compared TF motifs in accessible footprints at enhancers, promoters and gene bodies with or without the H3K18La peaks and detected enrichment of hundreds of TF binding motifs at H3K18La loci (**Figure 7A**). Enhancers and promoters with H3K18La in accessible regions present only 100-200 motifs from different TFs compared to an array of TFs with enriched motifs in gene bodies (**Figure 7A, B**), highlighting their distinct and specific regulatory roles. Notably, enriched TF motifs, especially in promoters and enhancers, exhibit a clear bias toward GC-rich sequences (**Figure 7C, Figure S3B-D**), consistent with a previous report ^29^. TFs whose motifs are enriched at H3K18La promoters and enhancers include several proteins of the KLF and the SP family as well as KMT2A, ZBTB14, TFDP1, VEZF1, PATZ1 and CTCF (**Figure 7C**, **Figure S4, Table S11-S19**). Interestingly, top motifs enriched at H3K18La gene bodies are for different TFs, such as SREBF2, MAZ, ZNF425, SALL4, HAND1 and 2 and SMAD4 (**Table S11-S19**). At specific loci, e.g., *Got 2* and *Eno1*, the accessible motifs at enhancers or promoters are concentrated within a single accessible footprint of each H3K18La peak (**Figure 7D-E**), whereas the various accessible motifs found in gene body of *Glis1* gene are distributed among several accessible footprints (**Figure 7F**). In retinal explants, a similar preference for accessible GC rich motifs is evident at H3K18La loci (**Figure S3F-G**). In retinal explants, similar to P28 retina, accessible H3K18La regions within gene bodies show a variety of motifs compared to enhancers or promoters (**Figures 7G, H**). Additional motifs are enriched in 25 mM compared to 5 mM glucose conditions for H3K18La, but not for H3K27Ac loci (**Figures 7G, H**).

**Figure 7:**
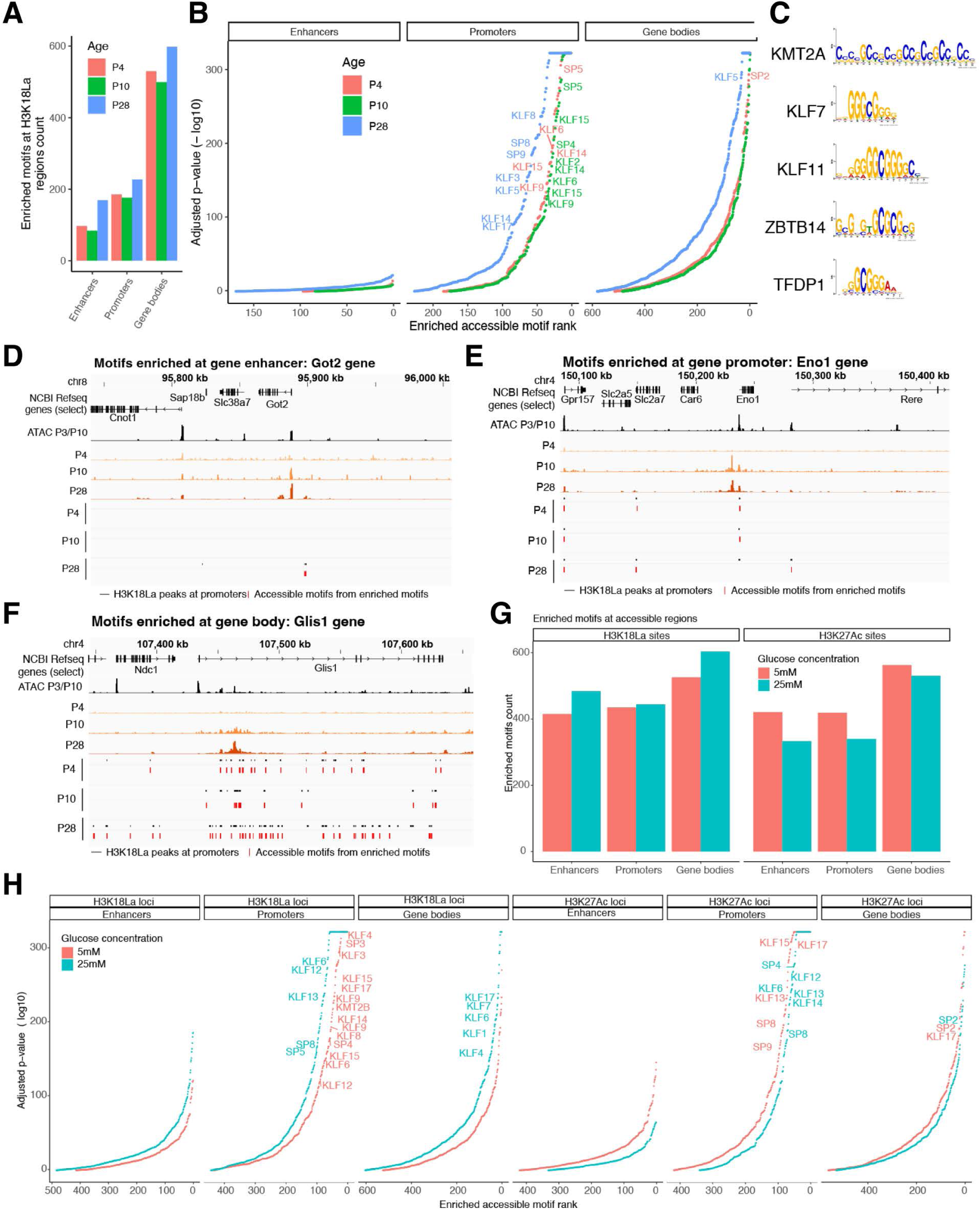
H3K18La promoters are enriched for GC rich binding motifs. (A) Number of enriched accessible motifs in H3K18La enhancers, promoters and gene bodies, at P4, P10 and P28. (B) Motif rank vs. p-value for motifs found at enhancers, promoters or gene bodies. (C) Top 5 accessible motifs enriched at H3K18La promoters at P28. (D-F) Examples of genes with H3K18La promoters containing accessible enriched motifs. Tracks represent H3K18La signal for P4, P10 and P28, H3K18La peaks (black) at either enhancers (D), promoters (E) or gene bodies (F) and accessible footprints containing enriched motifs at the given peaks (red) at P4, P10 and P28, and a selection of Refseq genes. One accessible footprint can contain multiple enriched motifs. (G) Number of enriched accessible motifs in H3K18La or H3K27Ac enhancers, promoters and gene bodies, for 5 or 25 mM glucose in explants. (H) Motif rank vs. p-value for motifs found at H3K18La or H3K27Ac enhancers, promoters or gene bodies for 5 or 25 mM glucose in explants. See also Figure S3 and S4

We conclude that H3K18La loci are enriched for GC rich accessible footprints. Presence of different motifs in enhancer/promoter versus gene body points to specific functions of H3K18La at distinct genomic locations.

## Discussion

Development, homeostasis and response to environment are coordinated by cooperative actions of genetic factors and epigenome modifications, which together facilitate precise patterns of gene expression ^47^. Cellular metabolism, and consequently availability of specific metabolites, can influence genome architecture. Notably, histone modifications play critical roles in establishing a permissive chromatin state for activating or repressing transcription ^8^. Unique physiological requirements of terminally differentiated retinal neurons present an attractive paradigm to explore the metabolism-epigenome nexus in development, aging and disease ^23,47,48^. In this report, we explore the direct role of aerobic glycolysis and lactate levels in modulating histone H3K18La and its contribution to gene expression. We show that enhanced lactate levels during retinal development and under high glucose conditions lead to changes in genome-wide profiles of H3K18La. The correlation of H3K18La to H3K27Ac and H3K4me3 profiles and to gene expression demonstrates a key role of histone lactylation in fine-tuning expression of both tissue-specific and ubiquitous genes.

Translation of genomic information is influenced by reversible epigenetic modifications of histones, which can help balance the transcriptional output under distinct cellular environments. Discovery of acyl marks on histones, such as propionyl, butyryl, crotonyl, and the more recent lactyl-CoA, has broadened the functional diversity of histone modifications ^27^. Histone acylations can mark genomic regions of embryonic tissues to generate distinct cell types ^30^. The profile of histone acylations in a specific cell type offers a snapshot of its metabolic status ^29,49^. Our studies provide strong evidence in support of gene regulation through histone KLa as a mechanism for functional maturation of retinal neurons during development and connects metabolic state, specifically lactate levels, to genetic controls. We also show that, unlike H3K27Ac, the regulation of gene expression by H3K18La is mediated by recruitment of select TFs. Our motif analysis reveals an enrichment for GC-rich accessible sequences at H3K18La peaks. The selection of a TF for binding to a specific region may be the result of relative dosage of each TF expressed in the retina as well as the competition between TFs with highly similar binding motifs. Nevertheless, H3K18La peaks in our study seems to be associated with zinc finger proteins, and the SP and KLF families. Our findings align with studies of genomic localization of H3K18La in various metabolic tissues, demonstrating its enrichment primarily at CGI-active gene promoters ^29^.

A recent study has demonstrated a remarkable association of lactate and histone acetylation (H3K27Ac) during early development using mouse eye organoids ^35^. Specifically, lactate was reported to inhibit Histone deacetylase 1, an enzyme that removes acetyl group and hence increased the enrichment of H3K27Ac near eye-field transcription factors within optic vesicle territory. We expand this energy-independent role of lactate as a direct epigenetic modifier of histones by lactylation at lysine residues. The results presented here directly correlate changes in aerobic glycolysis and consequently lactate to chromatin state and gene expression in developing retina in vivo and in mature retinal explants by manipulating glucose concentration. Enrichment of H3K18La in open chromatin and co-localization with H3K27Ac at many (but not all) genomic loci clearly suggest its key role in controlling retinal gene expression during development in altered metabolic (glucose) contexts.

Time periods between P6 and P10 and before eye opening (at around P14) mark significant developmental milestones in mouse visual system, with major changes in photoreceptor morphogenesis ^50^ and visual cortex ^51^. Our results (see Figure 1) showing changes in energy homeostasis during these developmental periods are consistent with retinal morphogenesis and specific upregulation of metabolic, phototransduction and developmental genes ^41^. Augmentation of genome-wide H3K18La marks at specific loci suggests the involvement of glycolysis/lactate/H3K18La regulatory axis in fine tuning of gene expression in maturing retina and directly linking the metabolic demands and glycolytic activity to transcription programs. A similar Glis1-driven epigenome-metabolome-epigenome signaling axis activates somatic cell reprogramming by repressing somatic genes and concurrently activating glycolysis-related genes to enhance lactate levels and histone modifications at pluripotency genes^31^.

Intriguingly, high glucose in explant retina cultures results in augmented H3K18La marks around the TSS of several photoreceptor genes (such as *Impg1, Impg2, Rp1, Gucy2f*), suggesting a potential mechanism of how photoreceptors respond to changes in local environment. Further investigations are needed to explore energy-independent functions of lactate as well as H3K18La-mediated transcriptional regulation in response to circadian rhythms, diet changes as well as during aging. Changes in glucose availability and/or utilization with aging, as reported in rods ^52^, may have implications for wider epigenome modifications (including histone lactylation and acetylation) that make the retina vulnerable to disease. Our study thus present a framework to elucidate the complex interplay among metabolism, gene regulation, and environmental factors and for understanding multifactorial etiology of age-related retinal diseases such as age-related macular degeneration (AMD) and diabetic retinopathy.

In conclusion, we show that H3K18La contributes to remodeling of the local chromatin environment and transcription in response to changes in metabolic state of the retina. Dynamics of histone lactylation at specific cis-regulatory elements likely helps in fine-tuning gene expression patterns in the retina during development and under distinct cellular environments. The reversible nature of histone lactylation presents an opportunity for designing therapies as well as dietary paradigms as indicated by epidemiological studies ^53,54^.

### Limitations of the study

Our studies provide a direct link between aerobic glycolysis and gene expression via H3K18La-mediated changes in chromatin environment. However, we appreciate the limitations of the retinal explant cultures, which do not recapitulate RPE-retina glucose shuttle. In addition, CUT&Tag studies provide genomic occupancy of H3K18La in the whole retina. As technical methods advance, single cell studies may uncover a more in-depth functional understanding of H3K18La in retinal cell types and its dysregulation in aging and disease.

## RESOURCE AVAILABILITY

### Lead contact

Further information and requests for resources and reagents should be directed to and will be fulfilled by the lead contact, Anand Swaroop.

### Materials availability

All unique/stable reagents generated in this study are available with a completed Materials Transfer Agreement per NIH policy. Further information and requests for resources and materials should be directed to and will be fulfilled by the lead contact, Anand Swaroop.

### Data and code availability

- All datasets used from public resources or produced in this study are summarized in the key resources table. The next generation sequencing data generated in this study are available at the Gene Expression Omnibus (GEO; accession number GSE291677).
- Custom code used for data analysis is available in GitHub (https://github.com/NEI-NNRL/2024_Mouse_Retina_H3K18La).
- Any additional information required to reanalyze the data reported in this paper is available from the lead contact upon request.

## Acknowledgments

We thank members of Swaroop laboratory, especially Anupam Mondal, Zachary Batz and Nivedita Singh, for discussions. This work was supported by Intramural Research Program of the National Eye Institute (ZIAEY000450 and ZIAEY000546) and utilized the computational resources of the NIH HPC Biowulf cluster (https://hpc.nih.gov). We also thank the CSHL mass spectrometry shared resource, supported by the Cancer Center Support Grant 5P30CA045508.

## Author contributions

Conceptualization, M.G. and A.S.; Methodology and Investigation, M.G., X.L., K.J., A.K., M.A.E., J.N., R.N.F., L.C.; Mass-spectrometry, P.C., M.C.P.; Bioinformatic analysis and Visualization, M.J.B., C.M.; Data submission, M.J.B.; Writing - Original Draft, M.G., X.L., M.J.B., C.M., L.C., A.S.; Writing, Review & Editing, all authors; Supervision, Project administration, and Funding acquisition, A.S.

## MATERIALS

### Key Resources Table

| REAGENT or RESOURCE | SOURCE | IDENTIFIER |
| --- | --- | --- |
| **Antibodies** |  |  |
| Anti-L-Lactyl-Histone H3 (Lys18) Rabbit mAb (ChIP Grade) | PTM BIO | Cat# PTM-1427RM |
| Anti-L-Lactyllysine Rabbit mAb | PTM BIO | Cat# PTM-1401RM |
| Anti-L-Lactyl-Histone H4 (Lys12) Rabbit mAb | PTM BIO | Cat# PTM-1411RM |
| Anti-Histone H3 (acetyl Lys27) Rabbit (ChIP Grade) | Abcam | Cat# ab4729 |
| Anti-Histone H3 Rabbit | Abcam | Cat# ab1791 |
| Anti-Histone H3 (acetyl Lys27) Mouse Monoclonal Antibody | Thermo Fisher Scientific | Cat# 2D7B3 |
| Alexa Fluor 488 donkey anti-rabbit IgG | Thermo Fisher Scientific | Cat# A-21206; RRID: AB_2535792 |
| Alexa Fluor 488 donkey anti-goat IgG | Thermo Fisher Scientific | Cat# A11055 |
| Alexa Fluor 568 donkey anti-mouse IgG | Thermo Fisher Scientific | Cat# A10037 |
| Human NRL Antibody | R&D SYSTEMS | Cat# AF2945 |
| Histone H3K27me3 Mouse Monoclonal Antibody | ACTIVE MOTIF | Cat# 61018 |
| DAPI | Thermo Fisher Scientific | Cat# D1306 |
| **Chemicals, peptides, and recombinant proteins** |  |  |
| pA-MNase conjugated to protein A | Steven Henikoff Lab |  |
| TRIzol® | Invitrogen | Cat# 15596026 |
| Oligomycin | Sigma | Cat# 75351 |
| Rotenone | Sigma | Cat# R8875 |
| 2-Deoxy-D-glucose | Millipore Sigma | Cat# D8375 |
| Glucose | Sigma |  |
| Neurobasal™-A Medium, no D-glucose, no sodium pyruvate | Thermo Fisher | Cat# A2477501 |
| B-27™ Supplement (50X) | Gibco™ | Cat# 17504044 |
| GlutaMAX™ Supplement | Gibco™ | Cat# 35050061 |
| Penicillin-Streptomycin-Glutamine (100X) | Gibco™ | Cat# 10378016 |
| Propionic anhydride | Sigma | Cat# 240311 |
| Tissue-Tek® O.C.T. Compound | Sakura Finetek USA | NA |
| Fluoromount-G mounting medium | SouthernBiotech | Cat# 0100-01 |
| **Critical commercial assays** |  |  |
| Lactate-Glo™ Assay | Promega | Cat# J5021 |
| CUT&Tag-IT<sup>®</sup> Assay Kit, Anti-Rabbit | ACTIVE MOTIF | Cat# 53171 |
| Agilent Seahorse XF Glycolytic Rate Assay Kit | Agilent | Cat# 1003346 |
| Agilent Seahorse XF Cell Mito Stress Test Kits | Agilent | Cat# 103015 |
| **Deposited data – Submitted to Gene Expression Omnibus (GEO)** |  |  |
| H3K18La CUT&Tag; development | This paper | GSE291664 |
| H3K18La CUT&Tag; explant | This paper | GSE291663 |
| H3K18La RNA-seq; explant | This paper | GSE291661 |
| **Experimental models: Organisms/strains** |  |  |
| C57BL/6J | Jax laboratory | N/A |
| Nrlp-GFP | Akimoto et al., 2006<br>55 | N/A |
| **Software and algorithms** |  |  |
| Image Studio | Licor | https://www.licor.com/bio/image-studio-lite/ |
| Proteome Discoverer 2.4 | Thermo Fisher |  |
| Orbitrap Fusion<sup>™</sup> Lumos<sup>™</sup> Tribrid<sup>™</sup> Mass Spectrometer | Thermo Fisher | Cat#IQLAAEGAAPF ADBMBHQ |
| AME (MEME) | https://meme-suite.org/meme/ | v5.5.7 |
| GraphPad Prism | Version 10.3.1 |  |
| nf-core/cutandrun | Nf-co.re | 3.1 |
| R | www.r-project.org | 4.2.3 |
| Bioconductor | www.bioconductor.org | 3.16 |
| tidyverse | CRAN | 2.0.0 |
| ChIPpeakAnno | Bioconductor | 3.32.0 |
| ChIPseeker | Bioconductor | 1.34.1 |
| org.Mm.eg.db | Bioconductor | 3.16 |
| clusterProfiler | Bioconductor | 4.6.0 |
| csaw | Bioconductor | 1.32.0 |
| edgeR | Bioconductor | 3.40.2 |
| PCAtools | Bioconductor | 2.10.0 |
| ggalluvial | CRAN | 0.12.4 |
| UpSetR | CRAN | 1.4.0 |

## EXPERIMENTAL MODEL AND STUDY PARTICIPANT DETAILS

### Mouse models

All procedures involving mice were approved by the Animal Care and Use Committee (NEI-ASP#650). C57BL/6J mice expressing EGFP under the control of the *Nrl* promoter (Nrlp-EGFP mice) ^55^ were used to perform Seahorse assays in the retina. Wild type mice (C57BL/6J) obtained from Jax laboratory were used for all other experiments unless specified. Mice were kept in a 12 light/12 dark hour cycle and fed *ad libitum* at the NEI animal facility.

### Retina Isolation and Explant Culture

Wild-type, P28 C57BL/6J mice used for retinal explant cultures were euthanized by CO_2_ asphyxiation and freshly dissected in 1X Hank’s Balanced Salt Solution (HBSS). To avoid variability due to circadian effects, retinas were dissected 2 hours after lights are turned ON in the facility for all experiments. Extraocular tissue was trimmed off and the cornea and iris were carefully removed. Sclera along with the RPE was gently removed to get the clean retina tissue. The dissected retinas were incubated in Neurobasal-A media supplemented with 5 mM glucose/25 mM glucose or 5 mM glucose with different concentration of 2-DG (10 mM and 20 mM), 0.2% B27 supplement, 0.1% N2 supplement, 0.1% Glutamax and 1X penicillin/streptomycin (Thermofisher Scientific) with 5% CO2 at 37°C for 48 hours times as described ^17^. At the end of incubation period, the retinas were rinsed with prewarmed Krebs’ Ringers medium and were further incubated in 0.3 mL Krebs’ Ringers medium for 1hr in 37°C incubator. The supernatant and retinas were rapidly frozen separately at the end of the experiment for lactate assay and western blot respectively. For CUT&Tag and RNAseq, retinas were processed immediately.

## METHOD DETAILS

### Histone extraction

Nuclear fractions were prepared from flash-frozen whole retina tissues obtained from four biological replicate of C57BL/6J mice samples of different age groups (P2, P6, P10, P14, P28). In brief, 50-100mg of tissue was Dounce homogenized on ice in hypotonic lysis buffer (10 mM Tris-HCl, 10 mM NaCl, 3 mM MgCl_2_, and 0.1% NP-40 alternative) with histone deacetylase and protease inhibitors and 1X phosphatase inhibitor. Crude sub-cellular fractions were separated by differential centrifugation at 800 x g for 10 minutes at 4°C. The crude nuclear pellet was then washed twice with ice-cold 1X PBS prior to acid extraction of histones using 0.4N H_2_SO_4_. The histones proteins were then precipitated using trichloroacetic acid with 0.1% sodium deoxycholate, washed with 100% acetone followed by elution in ultrapure water.

### Label-free chemical derivatization and mass spectrometry

Histone extracts from adult retina (n=3, P28) tissue samples were subjected to propionic anhydride chemical derivatization to react primary amines, as described ^56^. This reaction prevents derivatized lysines to be substrates for trypsin, thus restricting proteolysis to arginine sites, and reduces the overall charge state of the corresponding peptides making them amenable to LC/MS analyses. After derivatization, proteins were digested with sequencing grade modified porcine trypsin (Promega) overnight at 37 °C. Tryptic peptides were further treated with propionic anhydride to label newly generated N-termini. Derivatized peptides were desalted by reversed phase solid phase extraction using Pierce C18 Spin columns. Peptides were loaded on a 30 cm x 75 µm ID column packed with ReproSil 1.9 µm C18 particles (Dr. Maischt) and resolved on a 5-35% acetonitrile gradient in water (0.1% formic acid) using a Thermo Scientific Easy nLC 1200. Eluting peptides were analyzed by an Orbitrap Fusion Lumos mass spectrometer (Thermo). The MS was set to collect 120,000 resolution precursor scans (m/z 380-2000 Th). Precursor ions were selected for HCD fragmentation at stepped 28,33,38% NCE in a data-dependent manner and spectra were collected in the orbitrap at 60,000 resolutions with first mass locked to 100 Th.

### Immunohistochemistry

Eyes from 6-, 10-, 14- and 28-day-old C57BL/6J mice were enucleated, and the whole eyes were fixed in 4% (w/v) paraformaldehyde in phosphate-buffered saline (PBS) for 1 hour at room temperature. After being washed in PBS, the eyes were cryoprotected by sequential incubation in 10% and 20% sucrose in PBS for 1 hour at room temperature, followed by overnight incubation in 30% sucrose in PBS at 4°C. After the cornea, lens, and vitreous body were removed, the eyecups were embedded in optimal cutting temperature (OCT) medium, cut vertically at 10 µm on a Leica CM1850 cryostat (Wetzlar, Germany), and stored at −80°C until further use. Retinal sections were washed in PBS and blocked in 5% normal donkey serum in 0.5% Triton X-100 dissolved in filtered PBS for 1 hour at room temperature. Tissue sections were then incubated overnight at 4°C with the following primary antibodies: rabbit monoclonal anti-H3K18La (1:100, PTM1406RM, PTM Bio), or rabbit monoclonal anti-L-lactyllysine (Pan-lactyl) (1:200, PTM-1401RM, PTM Bio), mouse monoclonal anti-H3K27Ac (1:100, ThermoFisher Scientific Invitrogen 2D7B3), mouse monoclonal anti-H3K27me3 (1:100, Active motif, 61018) or human anti-NRL (1:100 R&D Systems, AF2945). Following three washes in PBS, slides were incubated with Alexa Fluor 488 donkey anti-rabbit IgG (H+L), Alexa Fluor 488 donkey anti-goat IgG, Alexa Fluor 568 donkey anti-mouse IgG and 1 μg/ml of 4’,6-diamidino-2-phenylindole (DAPI) for 1 hour at room temperature. Sections were then washed three times in PBS and mounted using Fluoromount-G mounting medium (SouthernBiotech, Birmingham, AL). Images were acquired with a Leica TCS SP8 at 40X or 63X magnification.

### Seahorse assay

Extracellular acidification rate (ECAR, as a measure of glycolysis flux) and Oxygen consumption rate (OCR, as a measure of mitochondrial respiration rate) were assessed on different age groups of Nrl-GFP mice (P2, P4, P6, P10, P14, P28) using Seahorse XFe24 Analyzer (Agilent, Santa Clara, CA, United States). Freshly dissected *ex vivo* retinal punches were prepared on the same day of the assay as described ^57^. The assays were performed using Agilent Seahorse XFe24 Extracellular Flux Assay kits and Seahorse XF24 islet capture microplates, with the Seahorse DMEM medium (6 mM of glucose, 0.12 mM of pyruvate, and 0.5 mM of glutamine). Four drugs: Oligomycin (final concentration 5 uM), Rotenone/Antimycin A (final concentration 1 uM) and 2-DG (final concentration 50 mM) were used in the assay. Basal glycoECAR and basal mitoOCR were determined using the measurement at 36 min, right before addition of the first drug, with formulars described previously ^57^. Glyco ATP production rate and mito ATP production rate were calculated using Agilent’s ATP production rate calculation algorithm (https://www.agilent.com/cs/library/whitepaper/public/whitepaper-quantify-atp-production-rate-cell-analysis-5991-9303en-agilent.pdf).

### Immunoblotting

Histone proteins were separated using 4-15% SDS-PAGE and transferred onto polyvinylidene fluoride (PVDF) membranes (Millipore). Blotted membranes were blocked in Licor blocking buffer and incubated with primary antibodies at 4°C overnight anti-Pan lactyl (PTM1401, PTM Bio), anti-H3K18La (PTM1406RM, PTM Bio), anti-H4K12La (PTM101, PTM Bio), anti-Histone H3 (ab176842, Abcam). Membranes were then washed three times with TBS-T and incubated with secondary antibodies at room temperature for 1 hr. The immunoreactive products were detected using Licor Odyssey Imaging System. ImageStudio was used to perform quantitative analysis of western blot results as the ratio of the band intensities of target histone marks to the band intensities of reference proteins (H3).

### Lactate assay

Lactate assay was performed on different age groups of mice (P2, P6, P10, P14, P28) using Promega Lactate-Glo (Promega J5021) assay according to the manufacturer’s instructions. Retina samples was homogenized in 1ml homogenization buffer with Inactivation solution at an 8:1 ratio using mechanical homogenizer. Tissue homogenate was used for protein estimation (Pierce BCA, 23225). Tissue lysates were neutralized with 0.125ml neutralization Solution and incubated with 50 μl detection reagent. Luminescence was recorded after 1 hour incubation using Promega™ GloMax® Plate Reader. The luminescence signal is further normalized with total protein concentration. The normalized luminescence values were plotted as proportional to lactate in the sample.

### CUT&Tag assay

CUT&Tag protocol was performed using CUT&Tag-IT® Assay Kit – Tissue (Active Motif) with slight modifications. In brief, retinas were dissected at P4, P10 and P28 and cells were dissociated in a 5 ml polypropylene round-bottom tube with a papain dissociation protocol adapted from previous study ^58^. After dissociation the cells were washed using 1X wash buffer provided in the kit at 200g for 5 minutes. CUT&Tag was performed using 500,000 cells per experiment following kit protocol. Antibodies against H3K18La (Rabbit, PTM1406RM, PTM Bio, China), and H3K27Ac (Rabbit, cat.no. ab4729, Abcam, Cambridge, UK) were used at a concentration of 1:100 in 100 μl and pA-MNase conjugated to protein A (generous gift of Dr. Steven Henikoff, Howard Hughes Medical Institute, Washington, USA) used at a concentration of 700 ng/ml. Libraries were paired-end sequenced to 101 bases using the NextSeq 2000 platform (Illumina, San Diego, CA).

### RNA extraction and library preparation

Total RNA from explant retinae was extracted using TRIzol® (Invitrogen, Carlsbad, CA), treated with DNase and cleaned up using the MagMAX mirVana Total RNA Isolation Kit (Applied Biosystems, Foster City, CA) following the manufacturer’s instructions. Libraries were constructed with SMARTer Stranded Total RNA-Seq Kit v2 – Pico Input Mammalian (Takara Bio USA, Mountain View, CA) with 4 ng of RNA and 13 PCR cycles library amplification. Paired-end sequencing of 101 bases were obtained using the NextSeq 2000 (Illumina, San Diego, CA). RNA-seq analysis pipeline was employed as previously described ^41^.

## QUANTIFICATION AND STATISTICAL ANALYSIS

### CUT&Tag Analysis

H3K18La CUT&Tag fastq files were processed using nf-core/cutandrun v3.1 to GRCm38 assembly <u>10.5281/zenodo.5653535</u> ^59^. MACS peaks having a post facto qValue (-log10) > 6 and present in at least two biological replicates were used as consensus peaks for each experimental grouping. Further analysis was performed in R v4.2.3 (r-project.org). Annotation of consensus peaks was performed with ChipSeeker v1.34.1 ^60^ using Ensembl v102 annotation with promoter defined from −1000 bp to +500 bp from transcriptional start site (TSS). Enhancer regions are defined as being −10kb to −1kb from TSS. Gene enrichment analysis was performed using clusterProfiler v4.6.0 with Gene Ontology Biological Process database (2022-Sep12 release). Peak quantitation for differential binding analysis was performed on merged consensus peaks P4, P10, and P28 using CSAW v1.32.0 ^61^. Normalization was performed using windowCounts function in CSAW with 10,000 base windows. PCA analysis was performed using PCAtools in R. Differential peak analysis was performed using gene wise negative binomial generalized linear models with quasi-likelihood tests using edgeR v3.40.2 ^62^. Explant analysis PCA was performed using peak quantitation from a merged 25 mM H3K18La and H3K27Ac consensus peaks. Previously published retina data was used in comparative analysis of H3K27ac and H3K4me3 ChIP-seq marks ^38^ and NRL CUT&Run ^39^. Previously published H3K18La peaks from different tissues ^29^ were processed using the same pipeline as the retina data for comparison. Upset plot for tissues was created from UpSetR v1.4.0 from presence of H3K18La peak in the promoter of protein coding genes. Hierarchical clustering (Ward’s method) of the Euclidean distance matrix of shared H3K18La promoter peaks between tissues was generated from the phi coefficient for all pairwise comparisons.

### Motifs enrichments analysis

We identified enriched motifs at H3K18La overlapping promoters, enhancers or gene bodies at P4, P10, P28, explant 5 mM or explant 25 mM glucose over control data. For each age ∼ region type pairs (e.g., P4 ∼ promoters), we selected the control data by taking all regions for the same type not overlapping any H3K18La at this age (e.g. all promoters without H3K18La peak at P4). We called footprint regions on these lists of regions (peaks and controls) using rgt-hint foot printing ^63^ with the ATAC-Seq option, using mapped ATAC-seq reads (SRR5884808, SRR5884809, SRR5884805, SRR5884806) ^38^. We extracted the DNA sequences from all footprint regions using bedtools getfasta ^64^. Finally, we ran motif enrichment analysis using MEME tool AME ^65^ (v5.5.7), with default parameters on peaks vs. control footprints, using the motifs database HOCOMOCO v12 ^66^ for Human and Mouse.

