## Supplemental Figures and Legends for "Aerobic glycolysis and lactate regulate histone H3K18Lactylation occupancy to fine-tune gene expression in developing and mature retina"

#### Figure S1

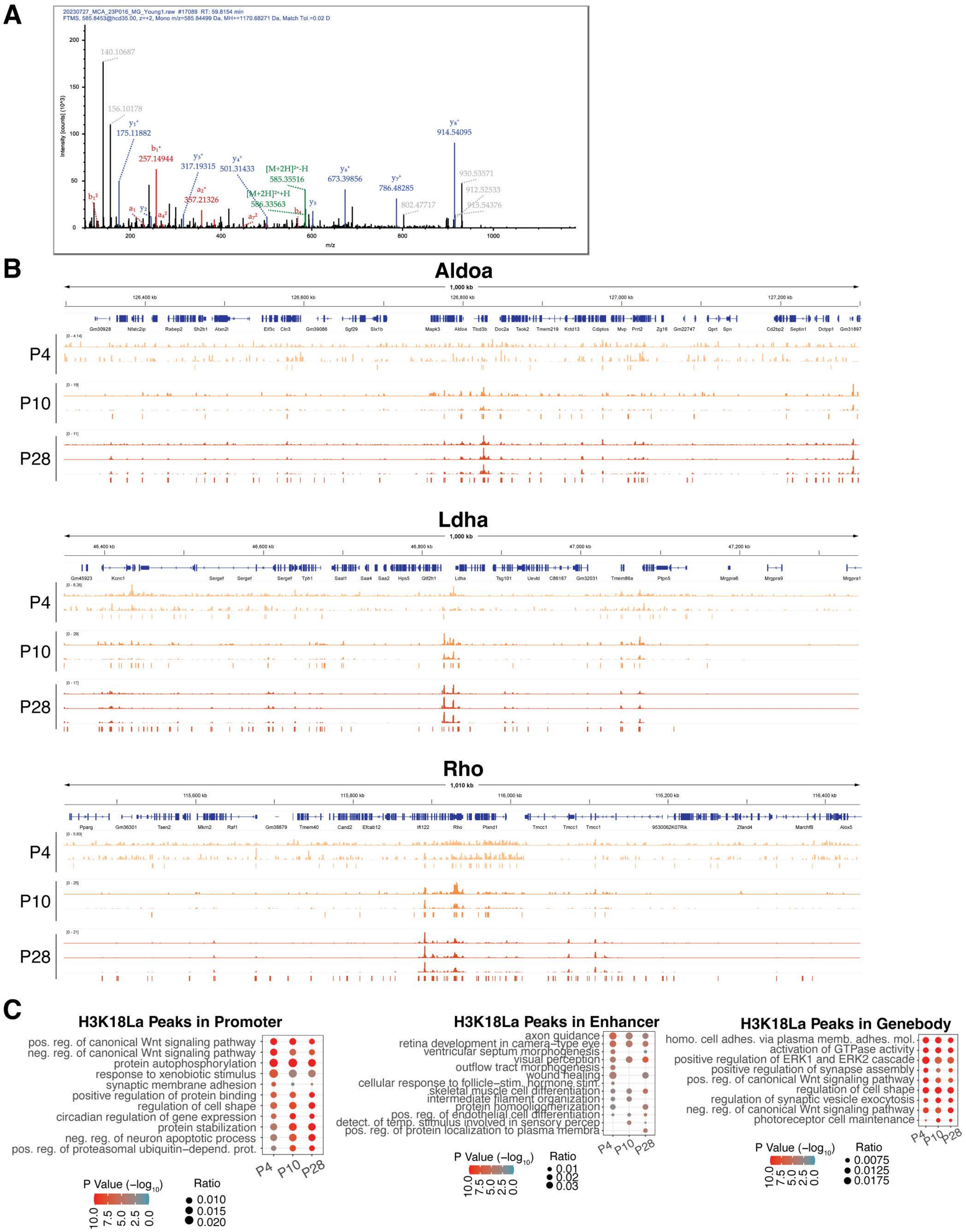
A
20230727_MCA_23P016_MG_Young1.raw #17089 RT: 59.8154 min
FTMS, 585.8453@hcd35.00, z=+2, Mono m/z=585.84499 Da, MH+=1170.68271 Da, Match Tol.=0.02 D
Intensity [counts] (10^3)
m/z
140.10687
156.10178
175.11882
257.14944
317.19315
501.31433
[M+2H]²⁺-H
585.35516
[M+2H]²⁺+H
586.33563
357.21326
673.39856
786.48285
914.54095
930.53571
912.52533
915.54376
802.47717
B
Aldoa
1,000 kb
Gm30928 Nfatc2ip Rabep2 Sh2b1 Atxn2l Eif3c Cln3 Gm39086 Sgf29 Slx1b Mapk3 Aldoa Tlcd3b Doc2a Taok2 Tmem219 Kctd13 Cdiptos Mvp Prrt2 Zg16 Gm22747 Qprt Spn Cd2bp2 Septin1 Dctpp1 Gm31897
P4
P10
P28
Ldha
1,000 kb
Gm45923 Kcnc1 Sergef Sergef Sergef Tph1 Saal1 Saa4 Saa2 Hps5 Gtf2h1 Ldha Tsg101 Uevld C86187 Gm32031 Tmem86a Ptpn5 Mrgpra6 Mrgpra9 Mrgpra1
P4
P10
P28
Rho
1,010 kb
Pparg Gm36301 Tsen2 Mkrn2 Raf1 Gm38879 Tmem40 Cand2 Efcab12 Ift122 Rho Plxnd1 Tmcc1 Tmcc1 Tmcc1 9530062K07Rik Zfand4 Marchf8 Alox5
P4
P10
P28
C
H3K18La Peaks in Promoter
pos. reg. of canonical Wnt signaling pathway
neg. reg. of canonical Wnt signaling pathway
protein autophosphorylation
response to xenobiotic stimulus
synaptic membrane adhesion
positive regulation of protein binding
regulation of cell shape
circadian regulation of gene expression
protein stabilization
neg. reg. of neuron apoptotic process
pos. reg. of proteasomal ubiquitin−depend. prot.
P4 P10 P28
P Value (−log10)
10.0 7.5 5.0 2.5 0.0
Ratio
0.010
0.015
0.020
H3K18La Peaks in Enhancer
axon guidance
retina development in camera−type eye
ventricular septum morphogenesis
visual perception
outflow tract morphogenesis
wound healing
cellular response to follicle−stim. hormone stim.
skeletal muscle cell differentiation
intermediate filament organization
protein homooligomerization
pos. reg. of endothelial cell differentiation
detect. of temp. stimulus involved in sensory percep
pos. reg. of protein localization to plasma membra
P4 P10 P28
P Value (−log10)
10.0 7.5 5.0 2.5 0.0
Ratio
0.01
0.02
0.03
H3K18La Peaks in Genebody
homo. cell adhes. via plasma memb. adhes. mol.
activation of GTPase activity
positive regulation of ERK1 and ERK2 cascade
positive regulation of synapse assembly
pos. reg. of canonical Wnt signaling pathway
regulation of cell shape
regulation of synaptic vesicle exocytosis
neg. reg. of canonical Wnt signaling pathway
photoreceptor cell maintenance
P4 P10 P28
P Value (−log10)
10.0 7.5 5.0 2.5 0.0
Ratio
0.0075
0.0125
0.0175

**Figure S1. Mass spectrum H3.3 peptide and genome-wide profiling of H3K18La, related to Figure 2B and Figure 4, respectively**

(A) Annotated HCD Spectrum of Histone H3.3 peptide R.KQLATKAAR.K modified with 1×Propionyl [N-Term]; 1×Propionyl [K6]; 1×lactylation [K]; 1×Acrolein [K1]. Peaks in blue are fragment ions containing the C-terminus (y-ions) and peaks in red are fragment ions containing the N-terminus (b-ions).

(B) Genomic histogram traces for 1 megabase regions of H3K18La sample replicated at each timepoint for genes involved in glycolysis (*Aldoa* and *Ldha*) and the phototransduction cascade in rod photoreceptors (*Rho*). The histogram traces are group scaled for each individual timepoint. The bars under each timepoint histogram represent consensus peaks.

(C) GO Biological Process gene sets enriched for genes containing H3K18La in each timepoint in different defined gene regions.

### Figure S2

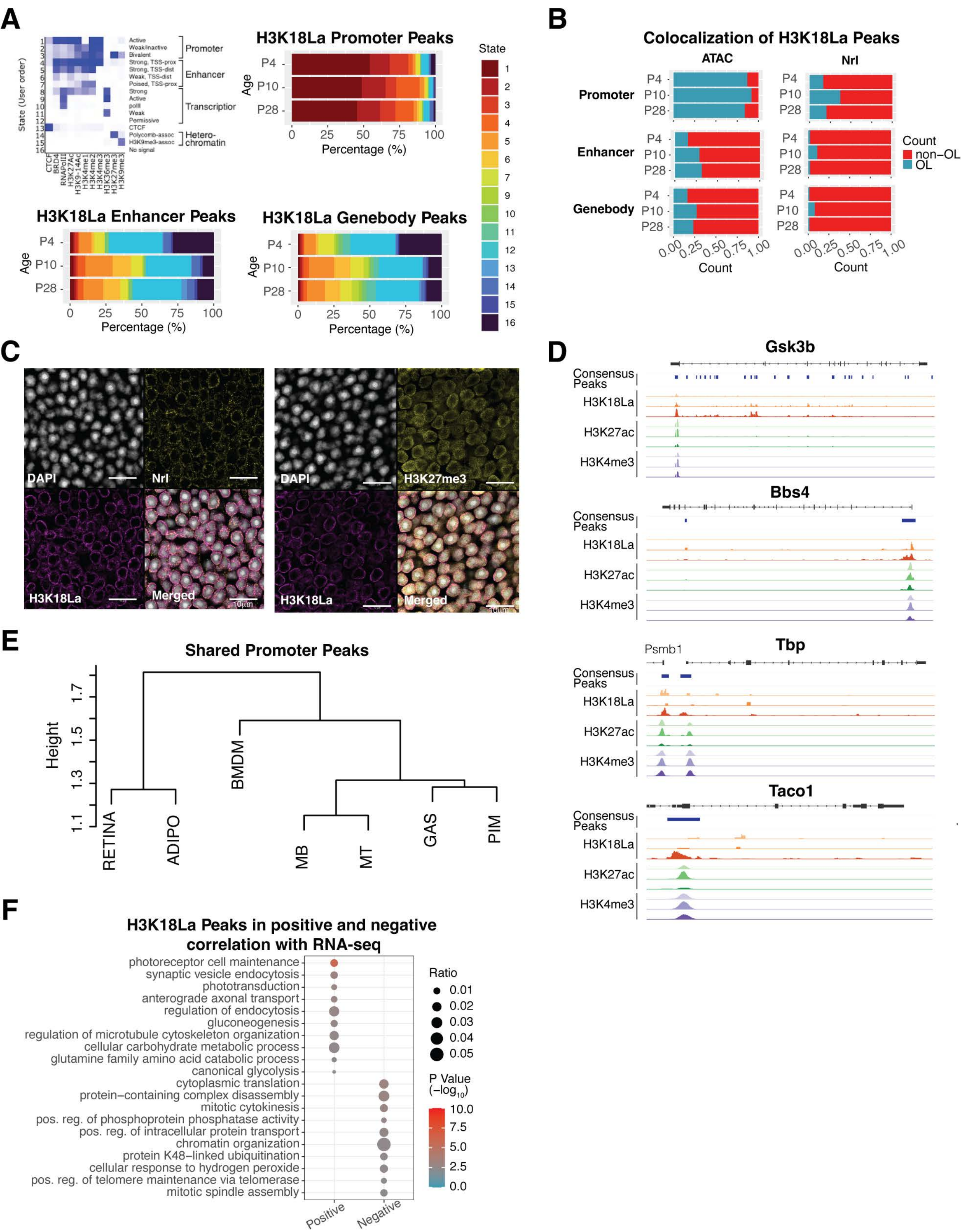
A
State (User order)
Active
Weak/inactive
Bivalent
Strong, TSS-prox
Strong, TSS-dist
Weak, TSS-dist
Poised, TSS-prox
Strong
Active
polII
Weak
Permissive
CTCF
Polycomb-assoc
H3K9me3-assoc
No signal
Promoter
Enhancer
Transcription
Hetero-chromatin
CTCF
BRD4
RNAPolII
H3K27Ac
H3K9-14Ac
H3K4me1
H3K4me2
H3K4me3
H3K36me3
H3K27me3
H3K9me3
H3K18La Promoter Peaks
H3K18La Enhancer Peaks
H3K18La Genebody Peaks
Age
P4
P10
P28
Percentage (%)
State
B
Colocalization of H3K18La Peaks
ATAC
Nrl
Promoter
Enhancer
Genebody
Count
non−OL
OL
C
DAPI
Nrl
H3K18La
Merged
H3K27me3
10μm
D
Gsk3b
Bbs4
Tbp
Psmb1
Taco1
Consensus Peaks
H3K18La
H3K27ac
H3K4me3
E
Shared Promoter Peaks
Height
RETINA
ADIPO
BMDM
MB
MT
GAS
PIM
F
H3K18La Peaks in positive and negative correlation with RNA-seq
photoreceptor cell maintenance
synaptic vesicle endocytosis
phototransduction
anterograde axonal transport
regulation of endocytosis
gluconeogenesis
regulation of microtubule cytoskeleton organization
cellular carbohydrate metabolic process
glutamine family amino acid catabolic process
canonical glycolysis
cytoplasmic translation
protein−containing complex disassembly
mitotic cytokinesis
pos. reg. of phosphoprotein phosphatase activity
pos. reg. of intracellular protein transport
chromatin organization
protein K48−linked ubiquitination
cellular response to hydrogen peroxide
pos. reg. of telomere maintenance via telomerase
mitotic spindle assembly
Positive
Negative
Ratio
P Value (−log10)

**Figure S2. H3K18La peak characterization and comparison, related to Figure 4**

(A) H3K18La peaks are overlapped with 16 chromatin states defined by ChromHMM from retina ChIP-seq data (Aldiri, et al., 2017) for each defined gene region and timepoint.

(B) Colocalization of H3K18La peaks with H3K27Ac (Aldiri, et al., 2017) bound regions.

(C) Confocal immunofluorescent images showing colocalization of Nrl and H3K27me3 with H3K18La in nuclear periphery of photoceptor cells. Scale bars, 10μm

(D) Genomic histogram traces of histone marks during development for representative selected genes found in Figure 4H. H3K18La marks at P4, P10, and P28 (Red), H3K27Ac (Aldiri, et al., 2017) at P3, P10, P21 (Green), and H3K4me3 at P3, P10, and P21 (Purple).

(E) Hierarchical clustering of shared H3K18La promoter peaks between retina and other tissues (Galle, et al., 2022).

(F) GO Biological Process gene sets enriched for protein coding genes containing H3K18La in the promoter which are positively or negatively correlated with RNA-seq expression during development.

Abbreviations: GAS, Gastrocnemius; MT, Post-mitotic end-state myotubes; PIM, Post-ischemia macrophages; MB, Myoblasts; ADIPO, Adipose tissues; BMDM, Bone marrow-derived macrophages.

### Figure S3

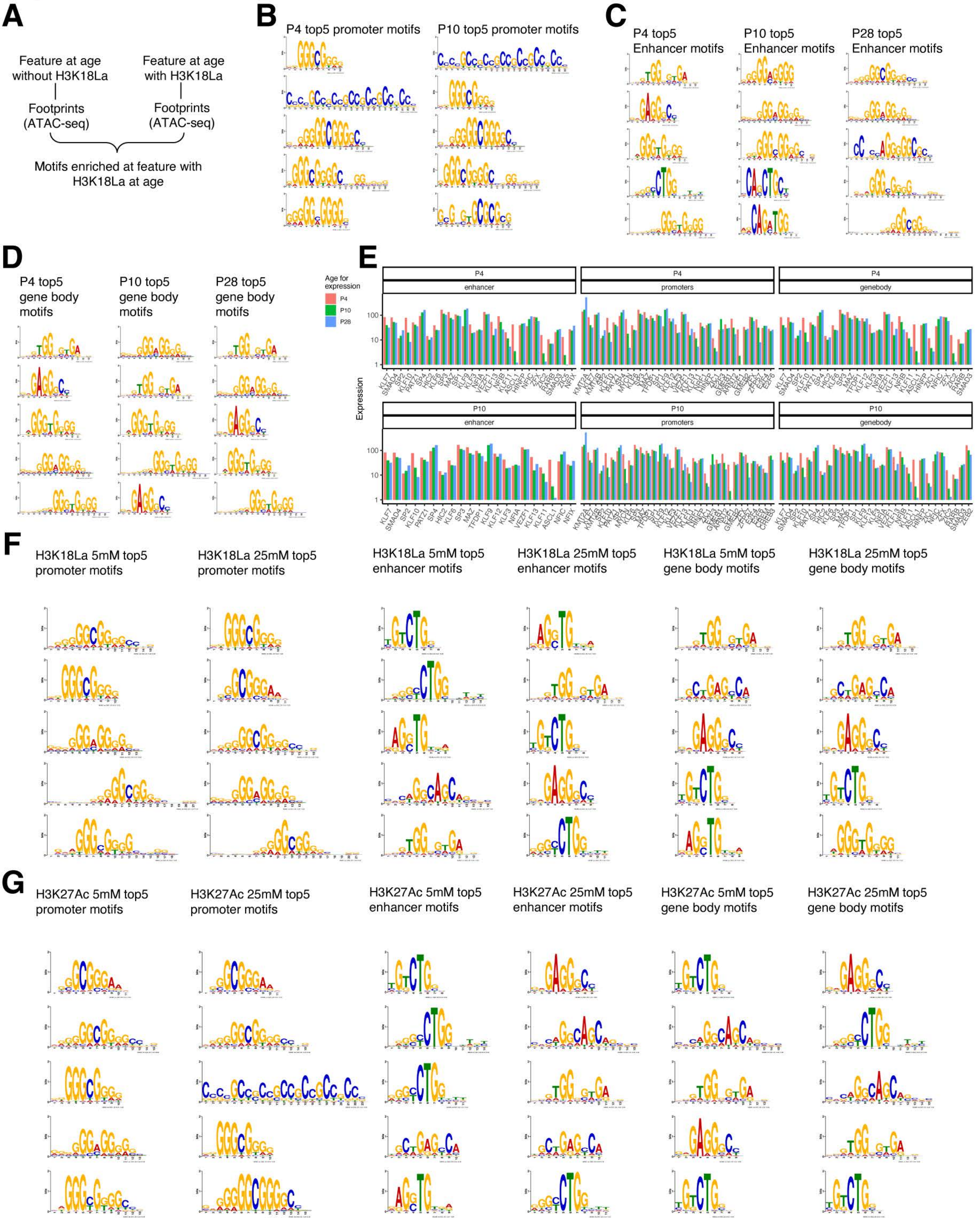
A
Feature at age without H3K18La
Feature at age with H3K18La
Footprints (ATAC-seq)
Footprints (ATAC-seq)
Motifs enriched at feature with H3K18La at age
B
P4 top5 promoter motifs
P10 top5 promoter motifs
C
P4 top5 Enhancer motifs
P10 top5 Enhancer motifs
P28 top5 Enhancer motifs
D
P4 top5 gene body motifs
P10 top5 gene body motifs
P28 top5 gene body motifs
E
Age for expression
P4
P10
P28
enhancer
promoters
genebody
Expression
F
H3K18La 5mM top5 promoter motifs
H3K18La 25mM top5 promoter motifs
H3K18La 5mM top5 enhancer motifs
H3K18La 25mM top5 enhancer motifs
H3K18La 5mM top5 gene body motifs
H3K18La 25mM top5 gene body motifs
G
H3K27Ac 5mM top5 promoter motifs
H3K27Ac 25mM top5 promoter motifs
H3K27Ac 5mM top5 enhancer motifs
H3K27Ac 25mM top5 enhancer motifs
H3K27Ac 5mM top5 gene body motifs
H3K27Ac 25mM top5 gene body motifs

**Figure S3. Enriched motifs during development for peaks containing H3K18La, related to Figure 7**

(A) Accessible motifs enrichment analysis pipeline.

(B) Top 5 accessible motifs enriched at H3K18La promoters at P4 (left) and P10 (right).

(C-D) Top 5 accessible motifs enriched at H3K18La enhancers (C) and gene bodies (D) at P4 (left) and P10 (center) and P28 (right).

(E) Expression levels of expressed TF from the 100 top enhancers, promoters or gene bodies motifs at P4 and P10.

(F-G) Top 5 accessible motifs enriched at H3K18La (F) or H3K27Ac (G) promoters, enhancers and gene bodies.

### Figure S4

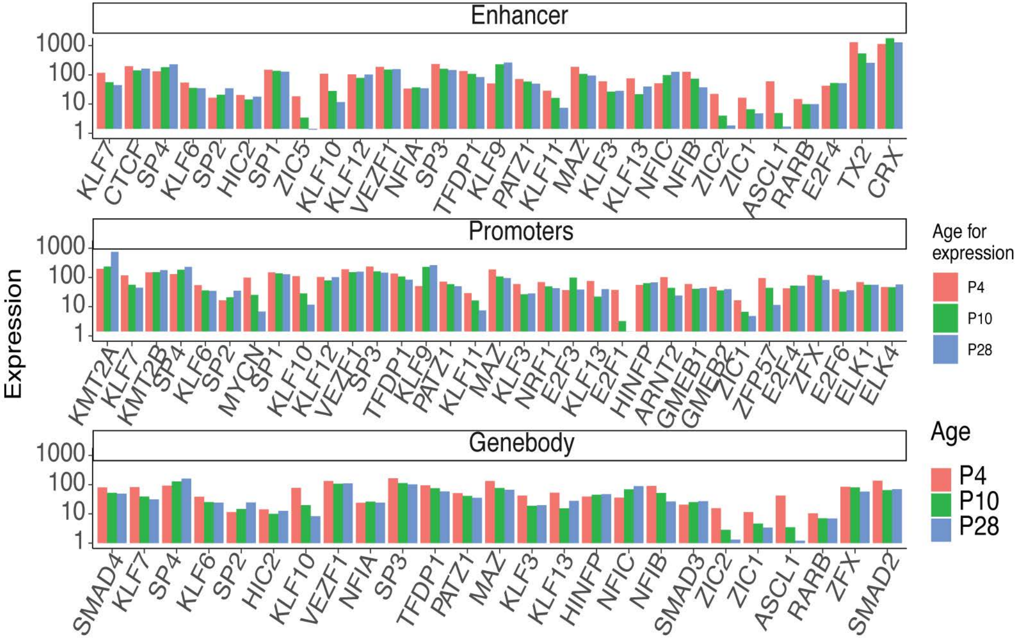
Enhancer
Promoters
Genebody
Expression
1000
100
10
1
KLF7 CTCF SP4 KLF6 SP2 HIC2 SP1 ZIC5 KLF10 KLF12 VEZF1 NFIA SP3 TFDP1 KLF9 PATZ1 KLF11 MAZ KLF3 KLF13 NFIC NFIB ZIC2 ZIC1 ASCL1 RARB E2F4 TX2 CRX
KMT2A KLF7 KMT2B SP4 KLF6 SP2 MYCN SP1 KLF10 KLF12 VEZF1 SP3 TFDP1 KLF9 PATZ1 KLF11 MAZ KLF3 NRF1 E2F3 KLF13 E2F1 HINFP ARNT2 GMEB1 GMEB2 ZIC1 ZFP57 E2F4 ZFX E2F6 ELK1 ELK4
SMAD4 KLF7 SP4 KLF6 SP2 HIC2 KLF10 VEZF1 NFIA SP3 TFDP1 PATZ1 MAZ KLF3 KLF13 HINFP NFIC NFIB SMAD3 ZIC2 ZIC1 ASCL1 RARB ZFX SMAD2
Age for expression
P4
P10
P28
Age
P4
P10
P28

**Figure S4. Enriched motifs in defined gene regions containing H3K18La, related to Figure 7**

Expression levels of expressed TF from the 100 top enhancers, promoters or gene bodies motifs at P28.

**Table S1. KLa peptide mass spectrometry results, related to Figure 2B**

Results from mass-spectrometry analysis, the identified lactylation sites on histone proteins when searched through the entire mouse database at all the different histone isoforms.

(XLSX)

**Table S2-S4. FGE results from H3K18La binding in promoter, enhancer, and gene body, related to Figure S1B**

Results from functional gene enrichment (FGE) analysis of Gene Ontology Biological Process at for genes with H3K18La peaks located in the promoter (**Table S2**), enhancer (**Table S3**), and gene body (**Table S4**) for each age investigated. GeneRatio indicates the number of genes identified in the pathway divided by the total number of genes at the age and peak location interrogated. BgRatio represents the number of genes in the ontology term divided by the total number of genes in the dataset. The ‘p.adjust‘ column indicates the Benjamini-Hochberg false discovery rate.

(XLSX)

**Table S5. FGE results from dynamic active histone mark (H3K4me3 and H3K27Ac) co-occupancy with H3K18La**, related to **Figure 4H**

Results from functional gene enrichment (FGE) analysis of Gene Ontology Biological Process for protein coding genes with H3K4me3, H3K27Ac, and H3K18La peak co-occupancy located in the promoter regions. GeneRatio indicates the number of genes identified in the pathway divided by the total number of genes at the age and peak location interrogated. BgRatio represents the number of genes in the ontology term divided by the total number of genes in the dataset. The ‘p.adjust‘ column indicates the Benjamini-Hochberg false discovery rate.

(XLSX)

**Table S6. FGE results from H3K18La peaks unique to retina, related to Figure 4J**

Results from functional gene enrichment (FGE) analysis of Gene Ontology Biological Process for protein coding genes with H3K18La peaks in promoter regions unique to retina. GeneRatio indicates the number of genes identified in the pathway divided by the total number of genes at the age and peak location interrogated. BgRatio represents the number of genes in the ontology term divided by the total number of genes in the dataset. The ‘p.adjust‘ column indicates the Benjamini-Hochberg false discovery rate.

(XLSX)

**Table S7. Differential binding analysis results, related to Figure 5B**

Results from differential binding analysis of H3K18La peaks at P10 versus P4 and P28 versus P4. Results fit a quasi-likelihood (QL) negative binomial generalized log-linear model (GLM) from the normalized count data. Log2 fold-change results are indicated in logFC.P10vsP4 and logFC.P28vsP4 columns. False discovery rate (FDR) indicates the Benjamini-Hochberg false discovery rate of the GLM QL fit. Columns P4_FPKM_mean, P10_FPKM_mean, and P28_FPKM_mean are the mean peak quantitation values in fragments per kilobase of peak per million (FPKM) mapped reads.

(XLSX)

**Table S8. FGE results from H3K18La peak quantitation correlated with RNA-seq expression, related to Figure S2F**

Results from functional gene enrichment (FGE) analysis of Gene Ontology Biological Process for quantitation of protein coding genes with H3K18La peaks in promoter regions during development which are positively and negatively correlated with RNA-seq expression from retina. GeneRatio indicates the number of genes identified in the pathway divided by the total number of genes at the age and peak location interrogated. BgRatio represents the number of genes in the ontology term divided by the total number of genes in the dataset. The ‘p.adjust‘ column indicates the Benjamini-Hochberg false discovery rate.

(XLSX)

**Table S9. FGE results from dynamic active histone mark (H3K4me3 and H3K27Ac) co-occupancy with H3K18La with positive correlation to RNA-seq, related to Figure 5E**

Results from functional gene enrichment (FGE) analysis of Gene Ontology Biological Process for protein coding genes with H3K4me3, H3K27Ac, and H3K18La peak co-occupancy located in the promoter regions. Dynamic occupancy is performed with genes showing positive correlation with RNA-seq gene expression during development. GeneRatio indicates the number of genes identified in the pathway divided by the total number of genes at the age and peak location interrogated. BgRatio represents the number of genes in the ontology term divided by the total number of genes in the dataset. The ‘p.adjust‘ column indicates the Benjamini-Hochberg false discovery rate.

(XLSX)

**Table S10. RNA-seq differential expression analysis results, related to Figure 6F**

Results from differential expression analysis of genes from retinal explants supplemented with 25mM versus 5mM glucose. Results fit a quasi-likelihood (QL) negative binomial generalized log-linear model (GLM) from the normalized count data. Log2 fold-change results are indicated in logFC column. False discovery rate (FDR) indicates the Benjamini-Hochberg false discovery rate of the GLM QL fit. Columns 5mM_CPM_mean and 25mM_CPM_mean are the mean gene expression values in gene counts per million (CPM) mapped reads.

(XLSX)

**Table S11-S19. TF motif analysis, related to Figure S4**

List of TF and their associated transcription measured by RNA-seq for P4 promoters (**Table S11**), P4 enhancer (**Table S12**), P4 gene body (**Table S13**), P10 promoter (**Table S14**), P10 enhancer (**Table S15**), P10 gene body (**Table S16**), P28 promoter (**Table S17**), P28 enhancer (**Table S18**), P28 gene body (**Table S19**)

(XLSX)
